# Normative and individual, non-normative intrinsic networks and the transition to impaired cognition

**DOI:** 10.1101/2024.02.20.581233

**Authors:** Qirui Zhang, Stacy Hudgins, Aaron F Struck, Ankeeta Ankeeta, Sam S. Javidi, Michael R Sperling, Bruce P Hermann, Joseph I. Tracy

**Affiliations:** Farber Institute for Neuroscience, Department of Neurology, Thomas Jefferson University, Philadelphia, Pennsylvania, USA; Department of Biomedical Engineering, Drexel University, Philadelphia, Pennsylvania, USA; Department of Neurology, University of Wisconsin School of Medicine and Public Health, Madison, Wisconsin, USA

**Author notes:** Correspondence to: Joseph I. Tracy Full address: Neuropsychology Division, Department of Neurology, Thomas Jefferson University/Vicky and Jack Farber Institute for Neuroscience, Health Professions Building, Department of Neurology, 4th floor, 901 Walnut Street, Philadelphia, PA 19107, USA.

**Keywords:** resting-state functional MRI, individualized brain networks, neuropsychology, seizure, verbal memory

## Abstract

Despite their temporal lobe pathology, a significant subgroup of temporal lobe epilepsy (TLE) patients are able to maintain normative cognitive functioning. Here, we identify TLE patients with intact versus impaired cognitive profiles, and interrogate for the presence of both normative and highly individual intrinsic connectivity networks(ICN) – all towards understanding the transition from impaired to intact neurocognitive status. We retrospectively investigated data from 88 TLE patients and matched 91 healthy controls with resting-state functional MRI. Functional MRI data were decomposed using independent component analysis to obtain individualized ICNs. Here, we calculated the degree of match between individualized ICNs and canonical ICNs (e.g., Yeo et.al 17 resting-state network) and divided each participant’s ICNs into normative or non-normative status based on the degree of match. We found that the individualized networks matched the canonical networks less well in the cognitively impaired compared to the cognitively intact TLE patients. The cognitively impaired patients showed significant abnormalities in the profiles of both normative and non-normative networks, whereas the intact patients showed abnormalities only in non-normative networks. At the same time, we found normative networks held a strong, positive association with the neuropsychological measures, with this association negative in non-normative networks. We were able to provide the initial data demonstrating that significant cognitive deficits are associated with the status of highly-individual ICNs, making clear that the transition from intact to impaired cognitive status is not simply the result of disruption to normative brain networks.

## 1. Introduction

Temporal lobe epilepsy (TLE) is associated with multiple domains of cognitive impairment, most commonly verbal memory.(Baxendale et al., 2010; Helmstaedter & Elger, 2009; Saling, 2009; Tai et al., 2018) Initially, the functional neuroanatomy of verbal memory deficits and other cognitive impairments such as executive function, language and sensorimotor skills was studied and explicated through a focus on regional brain functions, emphasizing specific one-to-one function (cognitive)/structural relationships.(Bell et al., 2011) More recent approaches have taken a network perspective. For example, models now emphasize that the structural damage to the hippocampus alters and degrades a broader, distributed memory network,(Caciagli et al., 2017; Lopez et al., 2022; Zhang et al., 2017) such that the sum of the disturbed information in the network leads to cognitive and behavioral dysfunction.(Bell et al., 2011) The implication is that the cognitive deficits in TLE patients are the result of disruptions to the organization and efficiency of the normative episodic memory system rather than damage to one specific brain structure.(Lariviere et al., 2020)

Subsequent functional magnetic resonance studies have validated this hypothesis, identifying the patterns of cognitive impairment that tend to be associated with alterations in well-defined (normative) resting-state or task-related networks,(Cataldi et al., 2013; Crow et al., 2023; Fadaie et al., 2021; He et al., 2018; He et al., 2022; Jokeit et al., 2001; Li et al., 2021; Rodriguez-Cruces et al., 2020; Szaflarski et al., 2017; Tracy et al., 2021) with each type of cognitive impairment displaying a unique set of normative network and connectome features that are perturbed by the underlying disease entity.(Ives-Deliperi & Butler, 2021) Missing from this network perspective has been the role played by highly individualized functional brain systems (non-normative networks) in cognitive function and dysfunction. Evidence has accrued to verify the presence of a large number of unique, individualized intrinsic networks in both healthy(Calhoun & de Lacy, 2017; Du et al., 2020; Laird et al., 2011) and diseased individuals.(Hinds et al., 2023; Maglanoc et al., 2020) For instance, in the setting of TLE, patient-specific networks have been identified, with evidence that the strength of such networks, in addition to the status of typical, normative intrinsic networks, can be predictive of outcomes following brain surgery.(Hinds et al., 2023) These data demonstrate the clinical value of assessing highly individualized networks, and raise the possibility that determining the network properties of these, in conjunction with normative intrinsic networks, may provide a broadened and improved view of the brain network damage associated with the progression of cognitive impairment in TLE.

Advances in our understanding of disease effects on cognition have indicated that the severity of cognitive impairment in TLE exists on a continuous spectrum.(Hermann et al., 2021; Tai et al., 2018; Zhao et al., 2014) In addition, there is now evidence that despite TLE pathology, a significant subgroup of patients are able to maintain normative cognitive functioning.(Coras et al., 2014; Tai et al., 2018) This adaptive brain response is usually attributed to cognitive reorganization, involving a change in the brain representation of cognitive functions. Extant models of cognitive reorganization in TLE emphasize the recruitment of compensatory, unaffected cognitive systems, housed in normative brain locations, to explain the maintenance of otherwise lost functionality.(Banjac et al., 2021; Chaitanya et al., 2020; McCormick et al., 2014) While imaging pathologic samples has demonstrated that our understanding of cognitive deficits is informed by disease impact on the regions and networks known to implement cognitive functions in healthy normals (for examples related to episodic memory, hippocampal integrity, and normative memory networks see(Coras et al., 2014; Lee et al., 2022)), we believe that integrating this with an understanding of the network properties of atypical or highly individual cognitive networks will greatly inform about both the deleterious impact of neuropathology and the nature of adaptive cognitive reorganization.

Previous studies have shown that cognitive status in TLE can be characterized by several cognitive phenotypes, involving one or more groups with expected cognitive anomalies, but also a group with an unexpected, intact cognitive profile.(Elverman et al., 2019; Hermann et al., 2020; Hermann et al., 2021; Reyes et al., 2019; Struck et al., 2023) In this context, a direct comparison of intrinsic brain systems in TLE patients with distinct neurocognitive profiles could have significant implications. We argue that investigating TLE patients with “intact” versus “impaired” neurocognitive profiles, interrogating for the presence of macroscopic intrinsic connectivity networks (normative and highly individual), characterizing the organizational properties of each, and, lastly, testing for a spectrum shift in these networks when transitioning from “intact-to-impaired” cognitive status,(Tracy et al., 2021) will help us understand how cognitively “intact” TLE patients support their cognitive functioning while other TLE patients display lost functionality.

More specifically, we hypothesize that in addition to normative brain networks, changes in non-normative brain networks are widely involved in both cognitive integrity and impairment in TLE. We use resting-state functional magnetic resonance independent component analysis (ICA) to capture and quantify individual, specific intrinsic connectivity networks (ICNs). We then estimate the “normativity” of individual brain networks by calculating the degree of match between individual ICNs and the canonical brain networks. For these normative and non-normative brain networks we characterize their network profiles by computing a set of core ICN metrics (number of ICNs and their underlying structural support), and ICN-related functional connection measures (ICN strength and network efficiency). This approach allows us to comprehensively assess the organizational features of these distinct brain networks, as well as examine the spectral properties associated with the remodeling/transitioning of cognition from “intact” to cognitively “impaired” status in TLE.

## 2. Materials and methods

### 2.1. Study Characteristics and Neuropsychological Measures

A total of 100 patients with drug-resistant unilateral TLE (60 left-sided, 40 right-sided) were recruited from the Thomas Jefferson Comprehensive Epilepsy Center. All patients had a seizure onset zone in the temporal lobe, established through the use of scalp EEG, MRI, PET, and other clinical data that determined this lobe harbors the seizure focus.(Sperling, 1993; Sperling et al., 1989; Sperling et al., 1992) We included pathology beyond classic hippocampal sclerosis to broaden the applicability of the study, as loss of connectivity or reserve is not dependent upon the presence of a gross macroscopic epileptogenic lesion.(Modi et al., 2021) Patients were excluded from the study for any of the following reasons: previous brain surgery; medical illness with central nervous system impact other than epilepsy; extratemporal or multifocal epilepsy; contraindications to MRI; or hospitalization for any Axis I disorder listed in the Diagnostic and Statistical Manual of Mental Disorders, V. Depressive disorders were allowed given the high comorbidity of depression and epilepsy. (Tracy et al., 2007)

Neurocognitive performance was assessed at the time of patient recruitment by a group of neuropsychological tests, as part of their standard clinical care and work up for potential surgical candidacy. The neuropsychological assessment included the following tests: general verbal intellectual abilities (Wechsler Adult Intelligence Scale(WAIS) III or IV -- vocabulary and -- similarities); language (Controlled oral word association test -- semantic fluency, -- letter fluency, and Boston naming test); executive functions (WAIS-III or IV -- matrix reasoning, -- digit span, Wisconsin card sort test-- perseverative responses, -- categories completed, Trail making test -- part B); mental efficiency and speed (Trail making test -- part A and WAIS- III or IV -- digit symbol coding); verbal learning and memory (Wechsler memory scale III or IV logical memory 1 and 2, California verbal learning test -- total learning and -- long delay free recall); visuospatial skills(WAIS-III or IV -- Block design, Rey–Osterrieth complex figure test – copy condition); psychomotor functioning (Grooved pegboard right and left hand). List, abbreviations, and citation of these test showed in Supplementary Materials and Methods.

At the same time, a total of 92 age-, and gender-matched healthy controls (HC) were recruited from the Thomas Jefferson University community. All controls were free of psychiatric or neurological disorders based on a health screening measure.

This study was approved by the Institutional Review Board for Research with Human Subjects at Thomas Jefferson University. All participants provided informed consent in writing.

### 2.2. K-means clustering analysis of neurocognitive performance

We used K-means clustering to perform cluster analysis on the neuropsychological data to obtain neurocognitive subtypes. No feature contained missing values greater than 10%, and we used mean interpolation for the missing values. Ultimately, 20 neuropsychological measures from 100 patients were included in the analysis. Based upon the K-means clustering results and verification of neuropsychological mean differences, we divided the patients into cognitively intact impaired group (see details in Results section).

### 2.3. Imaging Methods

All participants were scanned on a 3-T X-series Philips Achieva clinical MRI scanner (Amsterdam, Netherlands) using an 8-channel head coil. A 5-minute functional MRI scan involving a resting state condition and T1-weighted images were collected from all participants. During the resting-state condition, participants viewed a crosshair with no task requirements and were instructed to remain still throughout the scan, eyes open, and not fall asleep. The structural T1-weighted and functional data were pre-processed together in the fMRIPrep V22.1.1 pipeline.(Esteban et al., 2019) The eXtensible Connectivity Pipeline (XCP)(Ciric et al., 2018; Pruim et al., 2015; Satterthwaite et al., 2013) was used to post-process the outputs of fMRIPrep. The fMRI data were linearly detrended, denoised, filtered, smoothed, and finally aligned to 2mm MNI space. After head-motion estimation, we excluded 13 TLE patient with average head movement larger than 0.4 mm. Thus, 63 cognitively intact and 24 cognitively impaired patients will included in following image analysis. Details of the image data acquisition parameters, fMRIPrep and XCP processing details are described in the Supplementary Materials and Methods. After exclusion, fMRI head-motion (framewise displacement) of the three groups did not differ (cognitively intact, 0.16 ± 0.07 mm; cognitively impaired, 0.13 ± 0.06 mm; healthy controls, 0.15 ± 0.10 mm, F(192) = 0.90, p = 0.407), noting that the cognitively impaired group was slightly lower than the remaining two groups. Accordingly, this provides good evidence that head movement was not a potential cause of the observed brain network abnormalities in the cognitively impaired group.

### 2.4. ICN Metrics and ICN types

#### 2.4.1. Individualized ICN Construction

In the ICA, the fMRI dimensionality was estimated using multiple dimensions, d=20, 30, 40 on initial pass using FSL Melodic.(Smith et al., 2004) Optimized 30 ICNs were used for the main analysis based upon the mutual matching between individualized ICNs and canonical ICNs compared to other options (see the Yeo networks as an example, Supplementary Figure 1), and taking into account the experience of previous work.(Hinds et al., 2023; Laird, 2021; Smith et al., 2009; Wang & Li, 2015) This approach achieved a good balance between segmenting and combining regions to form coherent networks, ultimately yielding the optimal level of canonical ICN matching, while still providing a sufficient number of mismatched networks to allow analysis of non-normative, individualized ICN systems. The total variance explained by the 30 components is 88.8 ± 4.5%, average across samples.

#### 2.4.2. Matching individualized ICN with the canonical ICNs

To match a participant’s individualized ICNs (ICA components) to the established and functionally defined (i.e., canonical) intrinsic resting state systems, we used as canonical ICNs the 17 networks defined by Yeo et al. based on clustering analysis,(Yeo et al., 2011) and, subsequently for validation, the 10 networks reported by Smith et al. based on ICA.(Smith et al., 2009) The difference between these two templates is that the Yeo atlas is a mask with well-defined boundaries, dividing the cortex into 17 networks (Figure 1b). The Smith template, on the other hand, provides a whole-brain weighted distribution pattern of 10 canonical networks. We have adopted different strategies in matching individualized ICNs to these two atlas templates. For matching to the Yeo atlas, a threshold level of 0.5 (default in FSL) was used to exclude voxels that exceed the probability of being in the ‘background’ noise components in the individualized ICNs. The surviving voxels/regions were converted to binarized masks, and the dice coefficients were calculated separately for each individualized ICN with each of the 17 brain networks in the Yeo atlas. When matching Smith templates, we calculated spatial correlation coefficients between each individualized ICN and the 10 canonical ICNs. Best match for the canonical ICNs was defined as the individual component that matches best a particular canonical ICN, representing the match degree to each of the canonical ICNs. Best match for individualized ICNs was defined as the canonical ICN that matches best with an individualized ICN, representing the normativity of this person’s individualized ICNs. We then summarized the mean best matching value for canonical ICNs or individualized ICNs, representing the overall match of a subject to a canonical ICN (Figure 1c).

**Figure 1.**
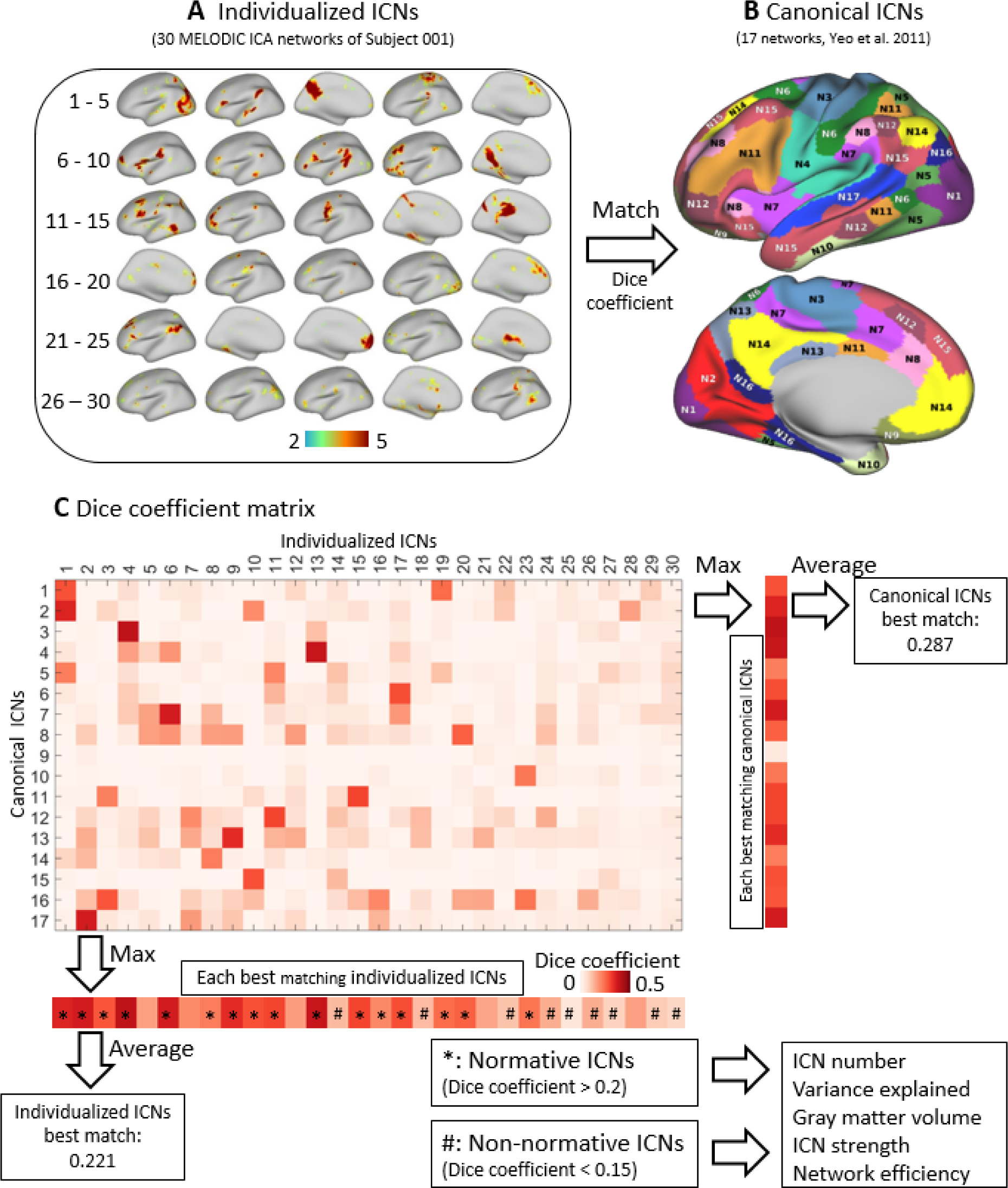
A case example of matching individualized ICN with canonical ICNs. **(a)** This a single participant data for illustrative purposes, 30 ICA components were used to represent individualized ICNs. **(b)** These networks were binarized; then dice coefficients were used to compute the degree of match with canonical ICNs (17 networks, Yeo et al. 37). **(c)** This cross-matching produced a matching matrix of 30 individualized ICNs ×17 networks. Based on this matrix we computed numerous matching properties as follows. Maximizing the matrix in the row direction obtained this individual’s best matching canonical ICNs, representing the match degree to each of the canonical ICNs for the analysis in Figure 3a. Maximizing the matrix in the column direction obtained the best matching individualized ICNs, representing the normativity of this person’s individualized ICNs. For all study participants, we categorized their individualized ICN’s as normative (*dice coefficient* > 0.2) or non-normative (*dice coefficient* < 0.15) based on thresholds. Lastly, 5 network measures of these two sets of ICNs were calculated and used in the analyses of Figure 3bc.

#### 2.4.3. Identifying and categorizing the level of ICN match

We identified the individualized ICNs that strongly matched to one of the canonical ICNs as normative ICNs. Conceptually, we considered individualized ICNs that demonstrated a “strong” match to one of the canonical ICNs to therefore represent that particular, cognitively normal, functional system, a system commonly and robustly found in healthy individuals and well established in the brain imaging literature.(Smith et al., 2009; Yeo et al., 2011) In contrast, we considered individualized ICNs demonstrating a “poor” match with the canonical ICNs to reflect coherent, systematic, but idiosyncratic variance in the resting state signal. These unmatched or poorly matched ICN components were highly individual, but cognitively and functionally undefinable as their spatial distribution did not resemble the 17 or 10 canonical networks. Accordingly, we regarded these ICNs as non-normative, noting that the number and nature of these ICNs varied by participant.

To identify the level of match to the canonical ICNs (i.e., to separate the normative from the non-normative ICNs), we used a “strong” match threshold to identify one of the person-specific, individual ICNs as a normative ICN. The individual ICNs with a low match value to each of the canonical ICNs (i.e., lower than a “poor” match threshold) were considered poorly matched to the canonicals and, therefore, non-normative. To define the thresholds for “strong” and “poor” matches, we used a threshold search of 0.05-0.5, with the optimal threshold defined as roughly equalizing the number of normative ICNs and the number of non-normative ICNs. We found that the best “strong” matching threshold for both the dice coefficient (Yeo template) and the spatial correlation coefficient (Smith template) was 0.2, while the best “poor” matching threshold was 0.15 for both (Supplementary Figure 2). Note, this threshold was the same used in a previous study (Hinds et al., 2023).

#### 2.4.4. ICN measures for the normative and non-normative ICNs

Based upon the normative and non-normative ICNs identified in each participant, separate for each ICN type, we computed the following metrics: number of components, total variance explained by the components, total ICN GM volume, total ICN connectivity strength, and overall efficiency (integration) among the components within each ICN type(Figure 1c). Briefly, for the ICN GM volume measure we calculated the GM volume underlying the regions in each ICN then summed the volume within each ICN type (normative, non-normative). ICN strength represented the strength of functional connectivity (edges) within each component, reflecting the level of intrinsic communication among the regions in a given ICN component. The overall efficiency between the components within each ICN type represented the level of functional integration (communication) among the components within ICN type (normative, non-normative). Detailed description of their calculations can be found in the Supplementary Materials and Methods.

### 2.5. Statistical Analyses

In our statistical analyses, we primarily utilized two-way mixed measures ANOVA to determine if the experimental groups (cognitively intact, cognitively impaired TLE patients, HCs) differed on the above ICN metrics and whether any such differences were a function of ICN type (normative, non-normative). Experimental group served as the between-subjects factor, and the network match or ICN measures served as the within-subject, mixed measures factor. The statistical significance threshold was set at *p* < 0.05 and the Tukey’s correction.

SmartPLS 4 software was used to perform PLS-SEM to test and specify the ICN measures most predictive of cognition performance within TLE patients. Among the advantages of the PLS-SEM model is the consideration of multicollinearity based on independent and dependent factors, which makes it superior to multiple regression methods. PLS-SEM was performed drawing a regression model between the ICN measures (*n* = 30) as a latent variable (independent factor) and neurocognitive performance (*n* = 20) as the target construct (dependent factor). Projections (outer loadings) of the ICN measures (latent variable) and neuropsychological measures (target construct) were calculated. The PLS-SEM utilized bootstrapping (5000 samples) to produce significance values for the outer loadings and the direct path coefficient between the ICN measure and neuropsychological latent factors. We also calculated the cross-correlations between ICN measures and neurocognitive performance using Pearson correlation, resulting a 30 × 20 correlation matrix.

### 2.6. Network subclasses in non-normative ICNs

Although the non-normative ICNs represented variable, person-specific networks, there may still exist commonalities and patterns among them. To identify the potential shared network patterns in the non-normative ICNs, we utilized clustering analysis. First, we defined the ICNs in each participant that matched neither the Yeo 17 (*dice coefficient* > 0.2) nor the Smith10 networks (*spatial correlation coefficients* > 0.2) as non-normative ICNs. We then pooled these non-normative ICNs (all such ICNs for all participants). Next, we used agglomerative hierarchical clustering to partition the non-normative ICNs into classes. Between-cluster distances were estimated using “Ward’s” minimum variance approach (Ward, 1963). As the non-normative ICNs contained a large number of specific networks, we needed to maximize our ability to differentiate between the various classes. Therefore, we set number of classes as 20, and then as a check on the stability of the results re-ran the clustering with number of classes as 40. By this method, each ICN network received a class label. As a measure of the frequency of each class label, we computed whether each participant owned, displayed each network class. Through chi-square tests, we then tested for between-group differences in the frequency of occurrence of each network class. The statistical significance threshold was set at 0.05 (FDR correction). Using the average ICN brain maps, we visually rendered the spatial patterns associated with each network class.

## 3. Results

### 3.1. Neurocognitive subtypes in TLE patients

We first analyzed cognitive covariance across subjects using Pearson correlation on 20 neuropsychological measures in 7 cognitive domains. Overall, we found these measures showed positive correlations in TLE patients and strong intra-cognitive domain covariance (Figure 2a). Through K-mean clustering, we found the group variance in the neuropsychological measures was best captured by a two-cluster solution. The cluster-wise Jaccard bootstrap proved the stability of these two cluster groups (Jaccard similarity = 0.894 and 0.810, for each group respectively). We found that cluster 2 (*n* = 28) was significantly lower than cluster 1 (*n* = 72) on all neuropsychological measures (all *p* < 0.01, FDR corrected). Therefore, we labelled cluster 1 as cognitively intact group and cluster 2 as cognitively impaired group. The most significant functional impairment was observed on the verbal learning and memory measures, with an average within-domain *t-statistic* mean of 6.62 (Figure 2b).

**Figure 2.**
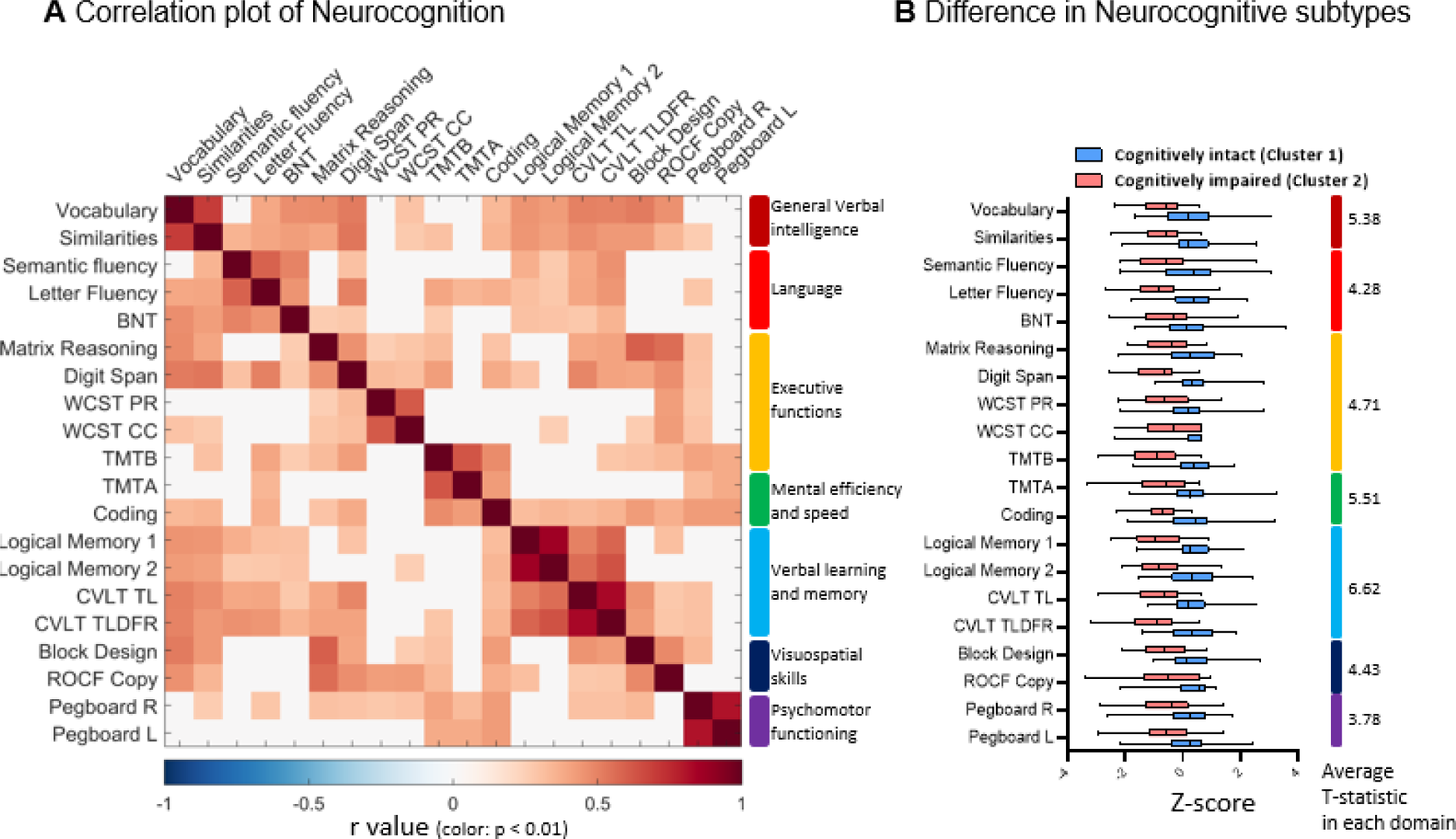
Neurocognitive subtypes of TLE patients. **(a)** Twenty neuropsychological measures in 7 cognitive domains included in analysis. Overall, these measures showed positive correlations in TLE patients and strong intra-cognitive domain covariance. **(b)** By K-mean clustering, we defined two patient groups: cluster 1(cognitively intact) and cluster 2 (cognitively impaired). The cognitively impaired group displayed significantly lower performances on all neuropsychological measures (all *p* < 0.01, FDR corrected). The most significant functional impairment was observed on the verbal learning and memory measures, with an average within-domain *t-statistic* mean of 6.62. Terms and abbreviations see Supplementary Materials and Methods.

The demographic and clinical characteristics of both groups of patients and HC are presented in Table 1. We did not find any between-group differences in age, sex, handedness, TLE laterality, and age at epilepsy onset, duration of epilepsy, temporal lobe pathology, or seizure type. Year of education was significantly higher in HC (*t*(190) = 4.25, p < 0.001), but there was no significant difference between cognitively intact and impaired patients (*t*(98) = 1.28, p = 0.203).

**Table 1.** Sample demographic and clinical characteristics.

| | Cognitively intact<br>(72) | Cognitively<br>impaired (28) | Healthy controls<br>(92) | F/t/$\chi^2$ | P |
| --- | --- | --- | --- | --- | --- |
| Age | 42.72 ± 13.65 | 40.25 ± 14.35 | 39.30 ± 12.66 | 1.363 | 0.258 |
| Sex(Male/Female) | 32/38 | 19/9 | 48/44 | 3.456 | 0.177 |
| Handedness(R/L) | 65/7 | 24/4 | 100/12 | 0.438 | 0.803 |
| Years of education | 14.76 ± 2.53 | 14.07 ± 2.12 | 16.16 ± 2.75 | 9.772 | <0.001 |
| TLE laterality(R/L) | 28/45 | 12/16 | N.A. | 0.171 | 0.678 |
| Age at epilepsy onset (year) | 25.86 ± 15.19 | 24.25 ± 16.82 | N.A. | 0.462 | 0.645 |
| Duration of epilepsy (year) | 17.33 ± 16.52 | 16.57 ± 14.96 | N.A. | 0.214 | 0.831 |
| Temporal pathology<br>(NB/HS/TE/TU/O) | 14/32/14/3/9 | 7/13/3/2/3 | N.A. | 1.628 | 0.803 |
| Seizure type |  |  | N.A. | 1.353 | 0.852 |
| FIAS | 1 | 1 |  |  |  |
| FAS | 13 | 6 |  |  |  |
| FAS + FIAS | 10 | 3 |  |  |  |
| FIAS + FBTCs | 27 | 12 |  |  |  |
| FAS + FIAS + FBTCs | 21 | 6 |  |  |  |
| Anti-epileptic drugs |  |  | N.A. |  |  |
| VGNC | 40 | 19 |  |  |  |
| VGCC | 4 | 4 |  |  |  |
| GABAa agonist | 3 | 3 |  |  |  |
| SV2a receptor mediated | 19 | 6 |  |  |  |
| CRMP2 receptor mediated | 32 | 8 |  |  |  |
| Multi-action | 8 | 8 |  |  |  |

Continuous variables are presented in mean ± SD.
Temporal pathology was diagnosed by neuro-radiologists based upon pre-surgical MRI scans: NB = normal brain; HS = hippocampal sclerosis; TE = temporal lobe encephalomalacia/gliosis, TU=tumor, O = Other MR abnormality (e.g., encephalocele, cavernoma).
Seizure types: FAS = focal onset aware; FIAS = focal onset impaired awareness; FBTCs = focal to bilateral tonic clonic.
Anti-epileptic drugs (Medication): VGNC = voltage-gated Na<sup>+</sup> channel blockage, e.g. phenytoin, carbamazepine, oxcarbazepine, lamotrigine (plus T Type Ca<sup>2+</sup> + channel blockage); VGCC = voltage-gated calcium channel, e.g. gabapentin; GABAa agonist, e.g. diazepam, clonazepam, clobazam, lorazepam, traxene, phenobarbital; SV2a receptor mediated, e.g. levetiracetam; CRMP2 receptor mediated, e.g. lacosamide (plus VGNC blockage); Multi-action: e.g. Na<sup>+</sup> + valproate (VGNC + GABAa agonist), topiramate (VGNC + GABAa agonist + AMPA/kainate receptor blockage + carbonic anhydrase inhibitor). Note, patients can be on more than one medication type. Patients may be on a medication regimen involving more than one drug type.
FD: framewise displacement.

### 3.2. Matching individualized ICNs with canonical ICNs

First, we performed individualized ICN decomposition of the resting-state functional MRI through ICA. Optimized 30 ICNs (detailed in Supplementary Figure 1) were used for the main analysis. By matching to each canonical ICN in Yeo 17 networks, we found significantly lower dice coefficients in the cognitively impaired patients by two-way mixed measures ANOVA (Group effect: *F*(16,2,176) = 6.445, *p* = 0.002, impaired vs. intact *p* = 0.028, intact vs. HC *p* = 0.429 and impaired vs. HC *p* = 0.001, Tukey’s corrected), with the largest decrease in limbic (temporopolar part) and dorsal attention (part A) networks (see Figure 3a, Supplementary Table 1).

**Figure 3.**
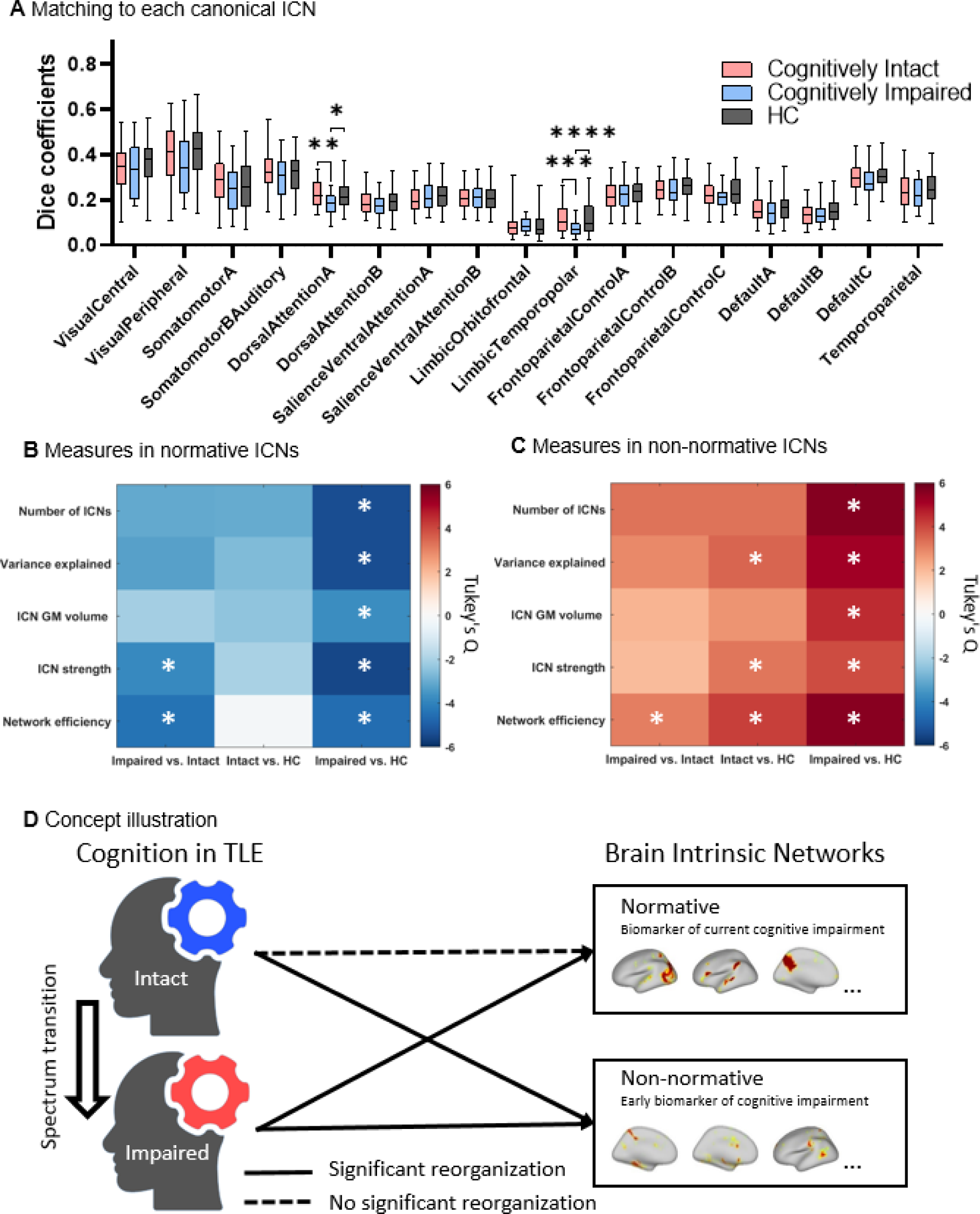
Matching individualized ICNs with canonical ICNs. **(a)** By matching to each canonical ICN in Yeo 17 atlas, significantly decreased dice coefficients in cognitively impaired patients were found by 3(groups) x 17(networks) two-way mixed measures ANOVA, especially in limbic (temporopolar part) and dorsal attention (part A) networks *: *p* < 0.05, **: *p* < 0.01, ***: *p* < 0.001, ****: *p* < 0.0001(Tukey’s corrected). **(b)** Post-hoc results in 3(groups) x 5(measures) two-way mixed measures ANOVA on ICN measures of normative and non-normative ICNs, with white asterisks indicating tests that were statistically significant (Tukey’s corrected). Color bar represents Tukey’s *Q* values. In general, measures of normative ICNs were significantly lower in cognitively impaired patients, but showed no difference relative to controls in cognitively intact patients. ICN strength and network efficiency showed a decrease compared to cognitively intact patients. **(c)** Same plot involving non-normative ICNs. In general measures of non-normative ICNs were significantly, abnormally increase in cognitively intact and impaired patients, with the cognitively impaired showing the highest network efficiency. (d) Concept illustration to summarize the result between cognition and type of brain networks.

This suggested that the individualized ICNs, overall, matched the normative network less well in cognitively impaired TLE patients. In addition, we found a strong correlation between the degree of match for the individualized ICNs and their variance explained across all participants (mean *r* = 0.645, standard deviation, *SD* = ± 0.188). This indirectly supports our hypothesis that the more normative the network, the more important it is in human cognitive function.

We then categorized each participant’s individualized ICNs by optimized thresholds (validation details in Supplementary Figure 2) as normative(dice coefficient > 0.2) or non-normative(dice coefficient < 0.15) ICNs based on the degree of match with the canonical ICNs, and calculated the following measures for each: number of ICNs, total ICN variance explained, ICN GM volume, ICN strength, and network efficiency. A two-way mixed measures ANOVA revealed, in general, a significant decrease in normative ICNs measures in the cognitively impaired patients, but no differences in the cognitively intact group. (Group effect: *F*(4,2,176) = 8.434, *p* < 0.001, impaired vs. intact *p* = 0.017, intact vs. HC *p* = 0.220 and impaired vs. HC *p* < 0.001, Tukey’s corrected, Figure 3b, Supplementary Table 2). In contrast, measures of non-normative ICNs were significantly increased in both cognitively intact and impaired patients, with this increase much larger in the impaired patients (Group effect: *F*(4,2,176) = 12.01 *p* < 0.001, Impaired vs. Intact *p* = 0.033, intact vs. HC *p* = 0.014 and impaired vs. HC *p* < 0.001, Tukey’s corrected, Figure 3c, Supplementary Table 3). These data indicated that the intact group has accrued an abnormal profile of non-normative ICNs, but only the impaired group showed a decreasing profile of the normative ICNs. We also validated the stability of the between-group differences under different matching thresholds using number of ICNs. The number of ICNs showed stable between-group differences with dice coefficients of 0.1-0.25 for the Yeo atlas, and spatial correlation coefficients of 0.05-0.35 for the Smith atlas (Supplementary Figure 2).

We noted that the ICN measures fell roughly into two categories. One, a set of core ICN metrics involving the number of ICNs, and their underlying functional (ICN variance explained) and structural (ICN GM volume) support. Two, ICN-related functional connectivity measures such as intra-network connection strength (ICN strength) and inter-network efficiency (network efficiency – an index of inter-network integration and communication). Utilizing post-hoc tests for each measure, we found that cognitively impaired patients differed significantly from controls on all ICN measures for both the normative and non-normative ICNs. Whereas the abnormalities relative to controls in the cognitively intact patients were only on the non-normative ICNs and were mainly related to the functional connectivity measures. (See Figure 3bc, Supplementary Figure 3, Supplementary Table 2, and Supplementary Table 3). We also plot a concept illustration to summarize the result between cognition and type of brain networks (Figure 3d).

We replicated these results with 20 and 40 individualized ICNs. The results indicated the findings remain stable across different numbers of ICN splits (see Supplementary Figure 4, Supplementary Figure 5). Moreover, by matching to each canonical ICN in the Smith 10 networks,(Smith et al., 2009) our results still showed the same trend, revealing significant differences between cognitively impaired patients and controls (Supplementary Figure 6). This further demonstrated the methodological stability of our frameworks for the ICN and neurocognitive groupings, as identical results were obtained with different atlases and matching algorithms. In addition to limbic and dorsal attention networks, in these various validation analyses the cognitively impaired patients also showed reduced matching in the default mode, visual, and fronto-parietal networks.

### 3.3. Relationship between ICN match and neurocognitive performance

Partial least squares regression analysis - structural equation modelling (PLS-SEM) was used to test and specify the ICN measures most predictive of cognitive performance within TLE patients. We found that modelling the ICN measures as a latent variable to explain the target construct (neurocognitive performance) produced significant results (model *R-square* = 0.155) with a *direct path coefficient* of 0.394 (*p* = 0.001, bootstrapping). With regard to projection to cognitive performance, measures from all the normative ICNs showed significant positive loadings, while all the non-normative ICN measures showed significant negative loadings (Figure 4a, Supplementary Table 4). The same pattern can be observed in cross-correlations between the ICN measures and cognitive performance (Figure 4bc).

**Figure 4.**
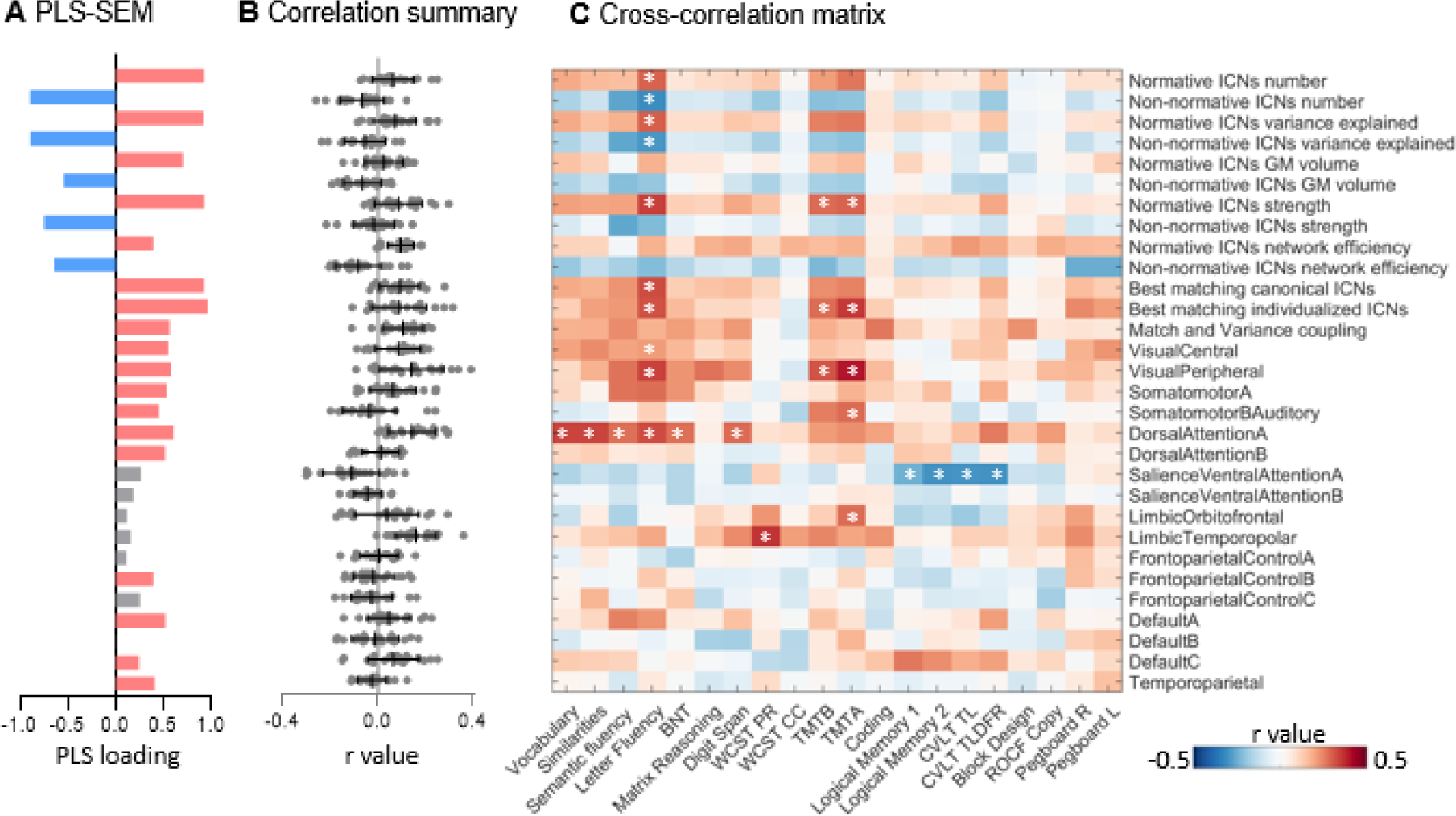
Relationship to neurocognitive performance. **(a)** PLS-SEM was used to test and specify the ICN measures most predictive of neurocognitive performance within TLE patients. The x-axis of the histogram is the loading of the ICN measures in the model. Red represents measures with significant positive loading, blue represents measures with significant negative loading, and gray represents measures with no significant loadings (bootstrapping corrected). All the normative ICN measures showed significant positive loadings, while all the non-normative ICN measures showed significant negative loadings. The degree of overall match and the match degree of specific canonical networks also showed positive loadings, with some measures being significant. **(b, c)** cross-correlations between ICN measures and neurocognitive performances. Correlation coefficients between each pair of measures are displayed in pseudo-color, with white asterisks representing correlations with p < 0.01. The correlation coefficient for each ICN measure with the 20 cognitive performances are summarized in **(b)**, each point represents the r value with neurocognitive performance. Note, the PLS-SEM loadings and cross-correlation r values displayed the same pattern on ICN measurements. Terms and abbreviations see Supplementary Materials and Methods.

These ICN/cognitive performance cross-correlations also explain specific test item correlations. For example, language function (letter fluency) and executive function (TMT part A and B) showed a relatively high correlation to ICN match measures. Also, the match degree of specific canonical networks produced a relatively high positive correlation to specific cognitive tests (dorsal attention network [part A] with general verbal intelligence and language function; limbic network [temporopolar part] with executive function; visual (peripheral) with language and executive functions, and a relatively high negative correlation between the salience ventral attention network [part A] to verbal learning and memory function (Figure 4c).

PLS-SEM results were likewise replicated by ICN matching to an alternative atlas of ICNs (Smith 10 networks(Smith et al., 2009)). The model explained slightly less variance in the neurocognitive performances construct (model *R-square*= 0.099), with a *direct path coefficient* of 0.316 (*p* = 0.219, bootstrapping). The coefficients of the outer loading variables was highly reproducible between the analyses of the Yeo and Smith networks, with a correlation of *r* (31) = 0.991 (*mean absolute difference* = 0.056), except for the match degree for each network, which was not comparable across the atlases (Supplementary Figure 7).

### 3.4. Network subclasses in non-normative ICNs

Finally, to understand the role of non-normative cognitive systems in impaired neurocognitive performance, we examined the spatial distribution of the non-normative ICNs relative to neurocognitive subtypes. We first performed a step of exploratory analysis, pooling together all 2692 non-normative ICNs of all the participants. We then utilized agglomerative hierarchical clustering to partition our non-normative ICNs into 20 subclasses (Figure 5a). Next, we computed whether each subject possessed ICNs belonging to each network class. Via chi-square tests, we found that, overall, cognitively impaired patients owned more ICNs in the cluster 1 and cluster 14 classes (Figure 5a, Supplementary Table 6). By topographically mapping the spatial distribution of these two subclasses on the brain, we found that Cluster 1 is predominantly a right lateralized language-related network, while cluster 14 is a thalamus-related network (Figure 5b). Also, through agglomerative hierarchical clustering, we categorized the non-normative ICNs into 40 categories, and found that it was still these two cluster classes that displayed reliable inter-group frequency differences (Supplementary Figure 8).

**Figure 5.**
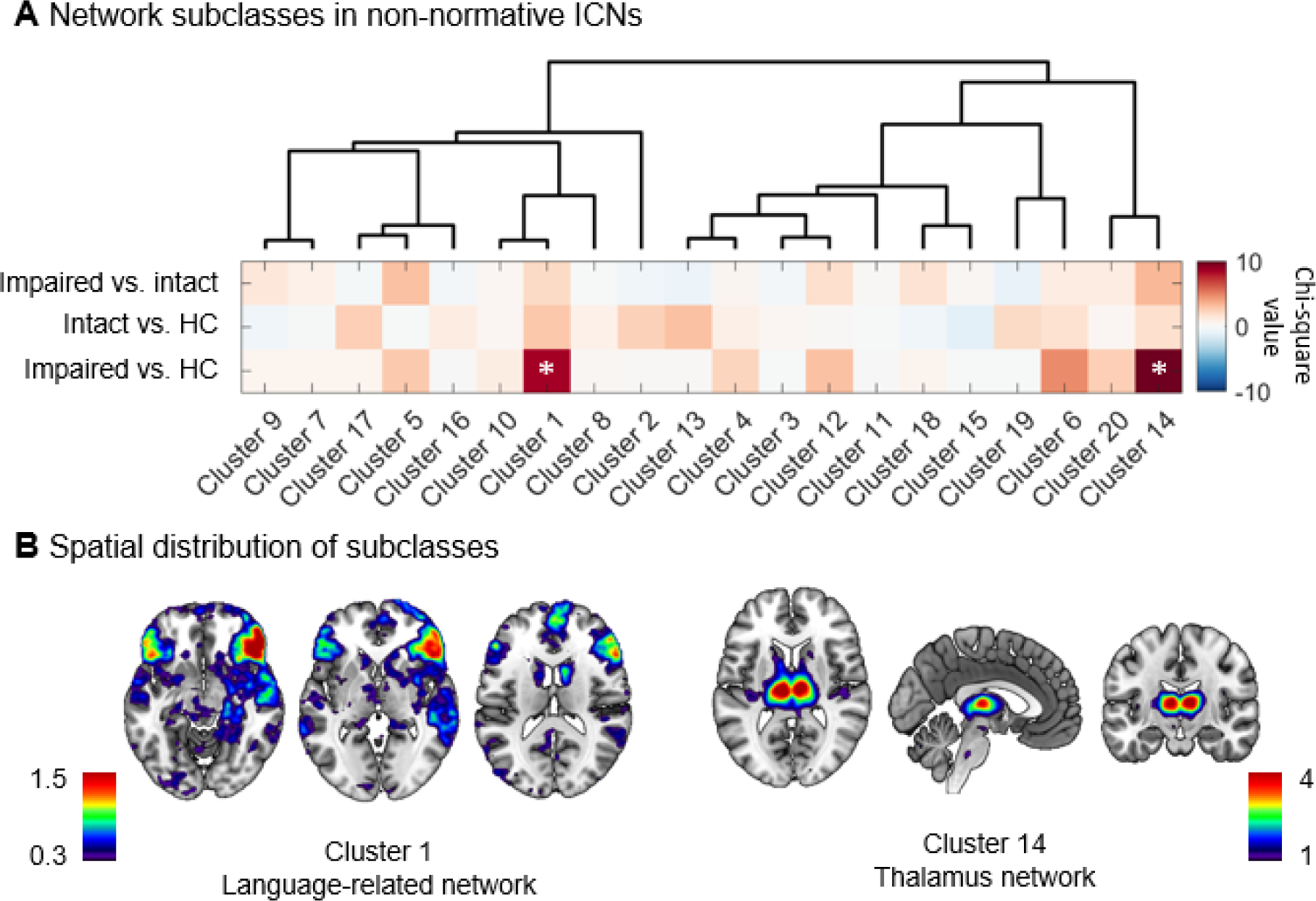
Network subclasses in non-normative ICNs. **(a)** Agglomerative hierarchical clustering partitioned the non-normative ICNs into 20 subclasses. Pseudo-colored plots show the chi-square value of differences in the frequency of occurrence of all subclasses across the study groups. Chi-square tests show the cognitively impaired patients possessed more ICNs in the cluster 1 and cluster 14 classes. **(b)** By mapping the brain spatial distribution of these two subclasses, we found that cluster 1 is predominantly a right lateralized language-related network, while cluster 14 is a thalamus-related network.

## 4. Discussion

By calculating the degree of match between individualized and canonical ICNs, we defined the normative or non-normative intrinsic brain networks of each participant. We found that the individualized ICNs matched the normative networks less well in the cognitively impaired TLE patients compared to the intact. This suggested that compared to the cognitively intact, the impaired patients possess larger sets of regionally specific coherent, systematic resting state networks that are both functionally distinct from the canonical networks and directly related to their weakened cognitive status. Both cognitively intact and impaired patients showed an abnormal increase in measures of non-normative brain networks, with this increase much larger in the cognitively impaired patients. This exacerbation was most notable with regard to the overall efficiency (inter-network integration) of these non-normative brain networks. However, only cognitively impaired patients showed declines in all profiles of normative brain networks.

Most notably, only cognitively impaired patients showed lower levels of match with the canonical brain networks. Particularly pronounced were abnormalities in multimodal networks including the dorsal attention network, default mode network, and frontoparietal network, all of which have been shown to be strongly associated with cognitive processes(Kaiser et al., 2015; Leech & Sharp, 2014; Menon, 2023; Shen et al., 2018) and cognitive deficits.(Ives-Deliperi & Butler, 2021; Kaiser et al., 2015) These reductions in network matching are, in general, atypical changes in network topology, changes that may explain why these networks could not provide cognitive compensation. We also found abnormalities for the cognitively impaired in the temporal pole part of the limbic system, which houses the core part of the temporal lobe epilepsy network over the cortex.(Caciagli et al., 2017; Lopez et al., 2022; Zhang et al., 2017)

In addition to quantifying the match with canonical networks, we also provide an overall description of normative brain network profiles in our two neurocognitive groups. Only the impaired group showed differences that go beyond these variations. These profiles were shown to have a positive impact on cognition (i.e., positive correlation) in our subsequent network analyses. This is consistent with data recognizing that network efficiency (integration) among the major networks is an important indicator of brain functional health.(Kim et al., 2017; Seguin et al., 2023; Vriend et al., 2020) This overall reduction in the number and efficiency of normative networks in cognitively impaired patients reflects dysfunction in normal cognitive processes and points to a reorganization of these normal cognitive systems. In contrast, cognitively intact patients did not display significant topological reorganization of the normative brain networks, noting that there was a certain decrease in the number of networks. Nonetheless, these patients still managed to have normal network efficiency, which may be the key to their support of intact neurocognitive function. This suggested that cognitive integrity is strongly linked to the topology of the well-established intrinsic networks.

More interesting, are our findings related to the profiles of non-normative brain networks. It is worth noting that, as expected, we found the normative ICN profiles held a strong, positive association with the neurocognitive measures, with this association negative for the non-normative ICNs. This makes clear the strong relation between cognitive impairment and non-normative ICNs. Overall, we found that cognitively impaired patients had obvious significant increases in all network measures, from the number of networks to network efficiency, whereas the profiles of cognitively intact patients also tended to be elevated, but only variance explained, ICN strength, and network efficiency were significantly different from controls. The increased efficiency among the non-normative networks in the intact patients suggested a tendency for these networks to integrate with each other, and it is possible that these highly individualized brain networks emerge to adaptively support cognitive activity despite also potentially being involved in TLE-related epileptic activity. There have been attempts in previous studies to try to isolate epileptic activity networks from individualized brain networks based on ICA by simultaneous EEG and functional magnetic resonance.(Ebrahimzadeh et al., 2021; Maziero et al., 2015; Rodionov et al., 2007) In any case, our data suggested that whatever role they play in cognition, at least in our cognitively intact patients, their alteration does not lead to significant cognitive impairment.

While the above discussion captures the general profiles of the individualized, non-normative brain networks, it is more intriguing to determine and specify the particular function these non-normative networks may be contributing in the patients with cognitive impairment. To do this, we performed a cluster analysis of these networks and found that there is some regional commonality across the non-normative networks, although, as expected, not as much as in the normative brain networks. Compared to controls, we found that the language and thalamus networks appeared more frequently in patients with cognitive impairment. The language network is generally a task-positive network,(Hickok & Poeppel, 2007) and its frequent appearance in the resting state may be due to a break in the balance between the task-positive and negative networks.(Liao et al., 2011; Yang et al., 2021) Alternatively, it may be that the patients’ language and memory functions are severely impaired and as part of the attempt to restore function these networks are strengthened during the resting state.(He et al., 2018) Indeed, previous has demonstrated that the strengthening of intrinsic networks during the resting state can work to enhance cognition at the time of actual performance.(Modi et al., 2021) The thalamus, on the other hand, serves as an important structure in temporal lobe epilepsy networks, potentially mediating focal-to-generalized seizures (He et al., 2020). The thalamus may also be an important, direct mediator of the spread of TLE structural damage originating in the temporal lobe,(Caciagli et al., 2017; Lopez et al., 2022; Zhang et al., 2017) and thalamic abnormalities are associated with poor surgical outcome.(He et al., 2017) Given the impaired status of these patients, we surmised that the communication systems identified by these particular non-normative networks are the most detrimental to overall cognitive functioning. An alternative possibility is that these non-normative networks reflect flawed, inchoate compensatory brain responses to TLE, responses that at least at the point of our assessment bore an association with dysfunction. The increased presence of these non-normative networks in the impaired patients, along with the reduced number of normative brain network, certainly points to topological reorganization of intrinsic systems in these patients.

Overall, despite the functional purpose of these non-normative networks remaining unconfirmed, a key implication of our data is that these non-normative networks play a role in cognitive dysfunction. Perhaps more surprisingly, our data for the intact patients showed that a select number of these non-normative networks, while not necessarily present (i.e., communicating) in every resting-state MRI session, are integral to supporting normal levels of cognitive functioning. Further investigation of their role in intact cognition needs to be pursued. We conclude that explanations of the cognitive deficits in impaired patients will come from an understanding of both their normative and individualized, non-normative intrinsic systems. While abnormalities in normative networks has been reported in the literature,(Cataldi et al., 2013; Crow et al., 2023; Fadaie et al., 2021; He et al., 2018; He et al., 2022; Jokeit et al., 2001; Li et al., 2021; Rodriguez-Cruces et al., 2020; Szaflarski et al., 2017; Tracy et al., 2021) by innovatively categorizing all individualized resting-state brain networks into normative and non-normative types, and calculating the profiles of each, we were able to provide the first data demonstrating that significant cognitive deficits are associated with the status of highly-individual ICNs, making clear that cognitive impairment is not simply the result of disruption to normative brain networks.

Importantly, our study reaffirmed that cognitive impairment in TLE involves multiple cognitive domains(Baxendale et al., 2010; Helmstaedter & Elger, 2009; Saling, 2009; Tai et al., 2018) as represented by the normative ICNs and their multivariate functionality. Through our cluster analysis, we showed that patients with cognitive impairments were broadly inferior to patients with intact cognitive performances, with verbal learning and memory the most severely impaired. However, our data also pointed to a potential relationship and transition mechanism between these two cognitive states. Given that even intact TLE patients displayed abnormalities compared to healthy controls in the profiles of non-normative networks, our data may imply that in TLE, changes to the integrity of these individualized brain systems is an early biomarker of cognitive impairment, signaling disruptions to intrinsic brain networks even before the disruptions to intrinsic normative networks becomes apparent. If this is the case, it argues that cognitive impairment lies on a spectrum, with the status of both the normative and highly-individualized, non-normative brain systems important to cognitive health and integrity. With this in mind, an additional implication of our data is that the breakdown of these two types of intrinsic systems may possess some order, with alterations in non-normative networks coming first, and the emergence of clear, observable functional deficits awaiting the disruption of normative intrinsic systems. In short, a key point in our data is that highly individualized networks were abnormal in TLE patients before they showed significant cognitive impairment. This spectrum perspective on the key cognitive phenotypes(Bell et al., 2011; Hermann et al., 2021; Struck et al., 2023) likely has implications for other disorders of cognitive impairment, perhaps even for normal aging. That is, highly individual, personal brain networks arising from an individual’s unique experiences and learning may possess an earlier vulnerability to brain disease compared to canonical brain networks. Accordingly, assessment of non-normative brain networks may be able to serve as early, more sensitive biomarkers for the predication and emergence of cognitive impairment in longitudinal studies.

Several aspects of our methodology are worth noting. Given our goal of identifying both canonical and non-canonical brain networks, the use of an individualized, data-driven approach is most appropriate. A difficult step in ICA studies is the identification or labelling of individualized brain networks typically done by comparing them with canonical brain network templates. This approach, however, discards a large number of components with potential physiological significance.(Calhoun & de Lacy, 2017; Du et al., 2020) Individualized brain network techniques based on a priori templates have been developed recently,(Cui et al., 2020; Cui et al., 2022; Kong et al., 2021; Nawaz et al., 2021; Wang et al., 2015) but these can only be used to identify individual topological variations in canonical brain networks. While the topology of an individual’s normative brain network is important to cognitive integrity, as our paper shows, our data more uniquely demonstrates the important role of non-normative brain networks. Before decomposing individual brain networks we used ICA-AROMA to remove head-movement related network components and regressed out potential covariates. This was done because the number and variance of physiological noise components (networks) varies from individual to individual, and a failure to suppress their presence could result in an uncontrollable proportion of physiological noise in our non-normative brain networks. In terms of the identification of normative and non-normative brain networks, different canonical brain network templates can produce different results. Accordingly, in this paper, two different atlases for canonical brain networks were utilized, each producing consistent results, demonstrating the stability of our strategy.

There are several limitations of this paper. First, for the non-normative brain networks, we only computed general profiles of their functionality. The specific functions and contributions of each and every type of network was not sufficiently explored. This would require case study reporting, or, alternatively large-scale samples to obtain robust estimates of the association between network organization/topologies and specific cognitive performances, going beyond the binary phenotypes of intact and impaired status that we analyzed. Second, we observed cognitive impairment at two points on the spectrum of TLE’s impact on functional brain networks, yet we only utilized cross-sectional data. Longitudinal approaches to map out the trajectory of intrinsic brain system alterations in association with aging and the potential progression or remission of TLE seizure activity is warranted. While our report contributes important data with regard to the potency of intrinsic resting network status in determining neurocognitive integrity, it will be important to test its added value compared to changes in other disease correlates, such as brain structure or the task-active networks most directly governing intact or impaired performances. Testing for correspondences (regional or otherwise) between these different influences on cognition would also be desirable, as no doubt, linkages and interactions exist between them. Lastly, limitations in our sample size may have contributed to our failure to find group differences on clinical variables (e.g. age of epilepsy onset, illness duration) or structural pathology(Coras et al., 2014; Lee et al., 2022), necessitating a replication of our reported effects with larger samples.

## 5. Conclusion

Cognitive impairment lies on a spectrum, with the status of both the normative and highly-individualized, non-normative brain systems important to cognitive health and integrity. With this in mind, our data can be seen as providing a demonstration of the substrates that transition the brain from intact to impaired cognitive status. More specifically, we show that the breakdown of these two types of intrinsic systems displays some order, with alterations in non-normative networks coming first, and the emergence of clear, observable functional deficits awaiting the disruption of normative intrinsic systems. Based on this, we conclude that non-normative brain networks may serve as early biological markers of forthcoming cognitive impairment.

## Contributors

Conceptualization: J.I.T., B.H, Q.Z.

Data curation: Q.Z., A.A., S.S.J.

Formal analysis: Q.Z., A.S.

Investigation: J.I.T., Q.Z., S.H., A.A., S.S.J., M.R.S

Methodology: Q.Z., J.I.T., B.H., A.S.

Visualization: Q.Z.

Supervision: J.I.T.

Writing—original draft: Q.Z.

Writing—review & editing: J.I.T.

## Acknowledgements

The authors are grateful and thank all the healthy individuals and patients with epilepsy, kept anonymous, who participated in this study. Work supported by Grant: JIT, NIH/NINDS, R01 NS112816-01.

## References

Banjac, S., Roger, E., Pichat, C., Cousin, E., Mosca, C., Lamalle, L., Krainik, A., Kahane, P., & Baciu, M. (2021). Reconfiguration dynamics of a language-and-memory network in healthy participants and patients with temporal lobe epilepsy. Neuroimage Clin, 31, 102702. 10.1016/j.nicl.2021.102702

Baxendale, S., Heaney, D., Thompson, P. J., & Duncan, J. S. (2010). Cognitive consequences of childhood-onset temporal lobe epilepsy across the adult lifespan. Neurology, 75(8), 705–711. 10.1212/WNL.0b013e3181eee3f0

Bell, B., Lin, J. J., Seidenberg, M., & Hermann, B. (2011). The neurobiology of cognitive disorders in temporal lobe epilepsy. Nat Rev Neurol, 7(3), 154–164. 10.1038/nrneurol.2011.3

Caciagli, L., Bernasconi, A., Wiebe, S., Koepp, M. J., Bernasconi, N., & Bernhardt, B. C. (2017). A meta-analysis on progressive atrophy in intractable temporal lobe epilepsy: Time is brain? Neurology, 89(5), 506–516. 10.1212/WNL.0000000000004176

Calhoun, V. D., & de Lacy, N. (2017). Ten Key Observations on the Analysis of Resting-state Functional MR Imaging Data Using Independent Component Analysis. Neuroimaging Clin N Am, 27(4), 561–579. 10.1016/j.nic.2017.06.012

Cataldi, M., Avoli, M., & de Villers-Sidani, E. (2013). Resting state networks in temporal lobe epilepsy. Epilepsia, 54(12), 2048–2059. 10.1111/epi.12400

Chaitanya, G., Hinds, W., Kragel, J., He, X., Sideman, N., Ezzyat, Y., Sperling, M. R., Sharan, A., & Tracy, J. I. (2020). Tonic Resting State Hubness Supports High Gamma Activity Defined Verbal Memory Encoding Network in Epilepsy. Neuroscience, 425, 194–216. 10.1016/j.neuroscience.2019.11.001

Ciric, R., Rosen, A. F. G., Erus, G., Cieslak, M., Adebimpe, A., Cook, P. A., Bassett, D. S., Davatzikos, C., Wolf, D. H., & Satterthwaite, T. D. (2018). Mitigating head motion artifact in functional connectivity MRI. Nat Protoc, 13(12), 2801–2826. 10.1038/s41596-018-0065-y

Coras, R., Pauli, E., Li, J., Schwarz, M., Rossler, K., Buchfelder, M., Hamer, H., Stefan, H., & Blumcke, I. (2014). Differential influence of hippocampal subfields to memory formation: insights from patients with temporal lobe epilepsy. Brain, 137(Pt 7), 1945–1957. 10.1093/brain/awu100

Crow, A. J. D., Thomas, A., Rao, Y., Beloor-Suresh, A., Weinstein, D., Hinds, W. A., & Tracy, J. I. (2023). Task-based functional magnetic resonance imaging prediction of postsurgical cognitive outcomes in temporal lobe epilepsy: A systematic review, meta-analysis, and new data. Epilepsia, 64(2), 266–283. 10.1111/epi.17475

Cui, Z., Li, H., Xia, C. H., Larsen, B., Adebimpe, A., Baum, G. L., Cieslak, M., Gur, R. E., Gur, R. C., Moore, T. M., Oathes, D. J., Alexander-Bloch, A. F., Raznahan, A., Roalf, D. R., Shinohara, R. T., Wolf, D. H., Davatzikos, C., Bassett, D. S., Fair, D. A., … Satterthwaite, T. D. (2020). Individual Variation in Functional Topography of Association Networks in Youth. Neuron, 106(2), 340–353 e348. 10.1016/j.neuron.2020.01.029

Cui, Z., Pines, A. R., Larsen, B., Sydnor, V. J., Li, H., Adebimpe, A., Alexander-Bloch, A. F., Bassett, D. S., Bertolero, M., Calkins, M. E., Davatzikos, C., Fair, D. A., Gur, R. C., Gur, R. E., Moore, T. M., Shanmugan, S., Shinohara, R. T., Vogel, J. W., Xia, C. H., … Satterthwaite, T. D. (2022). Linking Individual Differences in Personalized Functional Network Topography to Psychopathology in Youth. Biol Psychiatry, 92(12), 973–983. 10.1016/j.biopsych.2022.05.014

Du, Y., Fu, Z., Sui, J., Gao, S., Xing, Y., Lin, D., Salman, M., Abrol, A., Rahaman, M. A., Chen, J., Hong, L. E., Kochunov, P., Osuch, E. A., Calhoun, V. D., & Alzheimer’s Disease Neuroimaging, I. (2020). NeuroMark: An automated and adaptive ICA based pipeline to identify reproducible fMRI markers of brain disorders. Neuroimage Clin, 28, 102375. 10.1016/j.nicl.2020.102375

Ebrahimzadeh, E., Shams, M., Seraji, M., Sadjadi, S. M., Rajabion, L., & Soltanian-Zadeh, H. (2021). Localizing Epileptic Foci Using Simultaneous EEG-fMRI Recording: Template Component Cross-Correlation. Front Neurol, 12, 695997. 10.3389/fneur.2021.695997

Elverman, K. H., Resch, Z. J., Quasney, E. E., Sabsevitz, D. S., Binder, J. R., & Swanson, S. J. (2019). Temporal lobe epilepsy is associated with distinct cognitive phenotypes. Epilepsy Behav, 96, 61–68. 10.1016/j.yebeh.2019.04.015

Esteban, O., Markiewicz, C. J., Blair, R. W., Moodie, C. A., Isik, A. I., Erramuzpe, A., Kent, J. D., Goncalves, M., DuPre, E., Snyder, M., Oya, H., Ghosh, S. S., Wright, J., Durnez, J., Poldrack, R. A., & Gorgolewski, K. J. (2019). fMRIPrep: a robust preprocessing pipeline for functional MRI. Nat Methods, 16(1), 111–116. 10.1038/s41592-018-0235-4

Fadaie, F., Lee, H. M., Caldairou, B., Gill, R. S., Sziklas, V., Crane, J., Bernhardt, B. C., Hong, S. J., Bernasconi, A., & Bernasconi, N. (2021). Atypical functional connectome hierarchy impacts cognition in temporal lobe epilepsy. Epilepsia, 62(11), 2589–2603. 10.1111/epi.17032

He, X., Bassett, D. S., Chaitanya, G., Sperling, M. R., Kozlowski, L., & Tracy, J. I. (2018). Disrupted dynamic network reconfiguration of the language system in temporal lobe epilepsy. Brain, 141(5), 1375–1389. 10.1093/brain/awy042

He, X., Caciagli, L., Parkes, L., Stiso, J., Karrer, T. M., Kim, J. Z., Lu, Z., Menara, T., Pasqualetti, F., Sperling, M. R., Tracy, J. I., & Bassett, D. S. (2022). Uncovering the biological basis of control energy: Structural and metabolic correlates of energy inefficiency in temporal lobe epilepsy. Sci Adv, 8(45), eabn2293. 10.1126/sciadv.abn2293

He, X., Chaitanya, G., Asma, B., Caciagli, L., Bassett, D. S., Tracy, J. I., & Sperling, M. R. (2020). Disrupted basal ganglia-thalamocortical loops in focal to bilateral tonic-clonic seizures. Brain, 143(1), 175–190. 10.1093/brain/awz361

He, X., Doucet, G. E., Pustina, D., Sperling, M. R., Sharan, A. D., & Tracy, J. I. (2017). Presurgical thalamic “hubness” predicts surgical outcome in temporal lobe epilepsy. Neurology, 88(24), 2285–2293. 10.1212/WNL.0000000000004035

Helmstaedter, C., & Elger, C. E. (2009). Chronic temporal lobe epilepsy: a neurodevelopmental or progressively dementing disease? Brain, 132(Pt 10), 2822–2830. 10.1093/brain/awp182

Hermann, B., Conant, L. L., Cook, C. J., Hwang, G., Garcia-Ramos, C., Dabbs, K., Nair, V. A., Mathis, J., Bonet, C. N. R., Allen, L., Almane, D. N., Arkush, K., Birn, R., DeYoe, E. A., Felton, E., Maganti, R., Nencka, A., Raghavan, M., Shah, U., … Meyerand, M. E. (2020). Network, clinical and sociodemographic features of cognitive phenotypes in temporal lobe epilepsy. Neuroimage Clin, 27, 102341. 10.1016/j.nicl.2020.102341

Hermann, B. P., Struck, A. F., Busch, R. M., Reyes, A., Kaestner, E., & McDonald, C. R. (2021). Neurobehavioural comorbidities of epilepsy: towards a network-based precision taxonomy. Nat Rev Neurol, 17(12), 731–746. 10.1038/s41582-021-00555-z

Hickok, G., & Poeppel, D. (2007). The cortical organization of speech processing. Nat Rev Neurosci, 8(5), 393–402. 10.1038/nrn2113

Hinds, W., Modi, S., Ankeeta, A., Sperling, M. R., Pustina, D., & Tracy, J. I. (2023). Pre-surgical features of intrinsic brain networks predict single and joint epilepsy surgery outcomes. Neuroimage Clin, 38, 103387. 10.1016/j.nicl.2023.103387

Ives-Deliperi, V., & Butler, J. T. (2021). Mechanisms of cognitive impairment in temporal lobe epilepsy: A systematic review of resting-state functional connectivity studies. Epilepsy Behav, 115, 107686. 10.1016/j.yebeh.2020.107686

Jokeit, H., Okujava, M., & Woermann, F. G. (2001). Memory fMRI lateralizes temporal lobe epilepsy. Neurology, 57(10), 1786–1793. 10.1212/wnl.57.10.1786

Kaiser, R. H., Andrews-Hanna, J. R., Wager, T. D., & Pizzagalli, D. A. (2015). Large-Scale Network Dysfunction in Major Depressive Disorder: A Meta-analysis of Resting-State Functional Connectivity. JAMA Psychiatry, 72(6), 603–611. 10.1001/jamapsychiatry.2015.0071

Kim, J., Criaud, M., Cho, S. S., Diez-Cirarda, M., Mihaescu, A., Coakeley, S., Ghadery, C., Valli, M., Jacobs, M. F., Houle, S., & Strafella, A. P. (2017). Abnormal intrinsic brain functional network dynamics in Parkinson’s disease. Brain, 140(11), 2955–2967. 10.1093/brain/awx233

Kong, R., Li, J., Orban, C., Sabuncu, M. R., Liu, H., Schaefer, A., Sun, N., Zuo, X. N., Holmes, A. J., Eickhoff, S. B., & Yeo, B. T. T. (2021). Corrigendum to: Spatial Topography of Individual-Specific Cortical Networks Predicts Human Cognition, Personality and Emotion. Cereb Cortex, 31(8), 3974. 10.1093/cercor/bhab186

Laird, A. R. (2021). Large, open datasets for human connectomics research: Considerations for reproducible and responsible data use. Neuroimage, 244, 118579. 10.1016/j.neuroimage.2021.118579

Laird, A. R., Fox, P. M., Eickhoff, S. B., Turner, J. A., Ray, K. L., McKay, D. R., Glahn, D. C., Beckmann, C. F., Smith, S. M., & Fox, P. T. (2011). Behavioral interpretations of intrinsic connectivity networks. J Cogn Neurosci, 23(12), 4022–4037. 10.1162/jocn_a_00077

Lariviere, S., Rodriguez-Cruces, R., Royer, J., Caligiuri, M. E., Gambardella, A., Concha, L., Keller, S. S., Cendes, F., Yasuda, C., Bonilha, L., Gleichgerrcht, E., Focke, N. K., Domin, M., von Podewills, F., Langner, S., Rummel, C., Wiest, R., Martin, P., Kotikalapudi, R., … Bernhardt, B. C. (2020). Network-based atrophy modeling in the common epilepsies: A worldwide ENIGMA study. Sci Adv, 6(47). 10.1126/sciadv.abc6457

Lee, H. M., Fadaie, F., Gill, R., Caldairou, B., Sziklas, V., Crane, J., Hong, S. J., Bernhardt, B. C., Bernasconi, A., & Bernasconi, N. (2022). Decomposing MRI phenotypic heterogeneity in epilepsy: a step towards personalized classification. Brain, 145(3), 897–908. 10.1093/brain/awab425

Leech, R., & Sharp, D. J. (2014). The role of the posterior cingulate cortex in cognition and disease. Brain, 137(Pt 1), 12–32. 10.1093/brain/awt162

Li, Q., Tavakol, S., Royer, J., Lariviere, S., Vos De Wael, R., Park, B. Y., Paquola, C., Zeng, D., Caldairou, B., Bassett, D. S., Bernasconi, A., Bernasconi, N., Frauscher, B., Smallwood, J., Caciagli, L., Li, S., & Bernhardt, B. C. (2021). Atypical neural topographies underpin dysfunctional pattern separation in temporal lobe epilepsy. Brain, 144(8), 2486–2498. 10.1093/brain/awab121

Liao, W., Zhang, Z. Q., Pan, Z. Y., Mantini, D., Ding, J. R., Duan, X. J., Luo, C., Wang, Z. G., Tan, Q. F., Lu, G. M., & Chen, H. F. (2011). Default Mode Network Abnormalities in Mesial Temporal Lobe Epilepsy: A Study Combining fMRI and DTI [Article]. Hum Brain Mapp, 32(6), 883–895. 10.1002/hbm.21076

Lopez, S. M., Aksman, L. M., Oxtoby, N. P., Vos, S. B., Rao, J., Kaestner, E., Alhusaini, S., Alvim, M., Bender, B., Bernasconi, A., Bernasconi, N., Bernhardt, B., Bonilha, L., Caciagli, L., Caldairou, B., Caligiuri, M. E., Calvet, A., Cendes, F., Concha, L., … Group, E. N.-E. W. (2022). Event-based modeling in temporal lobe epilepsy demonstrates progressive atrophy from cross-sectional data. Epilepsia, 63(8), 2081–2095. 10.1111/epi.17316

Maglanoc, L. A., Kaufmann, T., Jonassen, R., Hilland, E., Beck, D., Landro, N. I., & Westlye, L. T. (2020). Multimodal fusion of structural and functional brain imaging in depression using linked independent component analysis. Hum Brain Mapp, 41(1), 241–255. 10.1002/hbm.24802

Maziero, D., Sturzbecher, M., Velasco, T. R., Rondinoni, C., Castellanos, A. L., Carmichael, D. W., & Salmon, C. E. (2015). A Comparison of Independent Component Analysis (ICA) of fMRI and Electrical Source Imaging (ESI) in Focal Epilepsy Reveals Misclassification Using a Classifier. Brain Topogr, 28(6), 813–831. 10.1007/s10548-015-0436-4

McCormick, C., Protzner, A. B., Barnett, A. J., Cohn, M., Valiante, T. A., & McAndrews, M. P. (2014). Linking DMN connectivity to episodic memory capacity: what can we learn from patients with medial temporal lobe damage? Neuroimage Clin, 5, 188–196. 10.1016/j.nicl.2014.05.008

Menon, V. (2023). 20 years of the default mode network: A review and synthesis. Neuron, 111(16), 2469–2487. 10.1016/j.neuron.2023.04.023

Modi, S., He, X., Chaudhary, K., Hinds, W., Crow, A., Beloor-Suresh, A., Sperling, M. R., & Tracy, J. I. (2021). Multiple-brain systems dynamically interact during tonic and phasic states to support language integrity in temporal lobe epilepsy. Neuroimage Clin, 32, 102861. 10.1016/j.nicl.2021.102861

Nawaz, U., Lee, I., Beermann, A., Eack, S., Keshavan, M., & Brady, R. (2021). Individual Variation in Functional Brain Network Topography is Linked to Schizophrenia Symptomatology. Schizophr Bull, 47(1), 180–188. 10.1093/schbul/sbaa088

Pruim, R. H. R., Mennes, M., van Rooij, D., Llera, A., Buitelaar, J. K., & Beckmann, C. F. (2015). ICA-AROMA: A robust ICA-based strategy for removing motion artifacts from fMRI data. Neuroimage, 112, 267–277. 10.1016/j.neuroimage.2015.02.064

Reyes, A., Kaestner, E., Bahrami, N., Balachandra, A., Hegde, M., Paul, B. M., Hermann, B., & McDonald, C. R. (2019). Cognitive phenotypes in temporal lobe epilepsy are associated with distinct patterns of white matter network abnormalities. Neurology, 92(17), e1957–e1968. 10.1212/WNL.0000000000007370

Rodionov, R., De Martino, F., Laufs, H., Carmichael, D. W., Formisano, E., Walker, M., Duncan, J. S., & Lemieux, L. (2007). Independent component analysis of interictal fMRI in focal epilepsy: comparison with general linear model-based EEG-correlated fMRI. Neuroimage, 38(3), 488–500. 10.1016/j.neuroimage.2007.08.003

Rodriguez-Cruces, R., Bernhardt, B. C., & Concha, L. (2020). Multidimensional associations between cognition and connectome organization in temporal lobe epilepsy. Neuroimage, 213, 116706. 10.1016/j.neuroimage.2020.116706

Saling, M. M. (2009). Verbal memory in mesial temporal lobe epilepsy: beyond material specificity. Brain, 132(Pt 3), 570–582. 10.1093/brain/awp012

Satterthwaite, T. D., Elliott, M. A., Gerraty, R. T., Ruparel, K., Loughead, J., Calkins, M. E., Eickhoff, S. B., Hakonarson, H., Gur, R. C., Gur, R. E., & Wolf, D. H. (2013). An improved framework for confound regression and filtering for control of motion artifact in the preprocessing of resting-state functional connectivity data [Article]. Neuroimage, 64, 240–256. 10.1016/j.neuroimage.2012.08.052

Seguin, C., Sporns, O., & Zalesky, A. (2023). Brain network communication: concepts, models and applications. Nat Rev Neurosci. 10.1038/s41583-023-00718-5

Shen, X., Cox, S. R., Adams, M. J., Howard, D. M., Lawrie, S. M., Ritchie, S. J., Bastin, M. E., Deary, I. J., McIntosh, A. M., & Whalley, H. C. (2018). Resting-State Connectivity and Its Association With Cognitive Performance, Educational Attainment, and Household Income in the UK Biobank. Biol Psychiatry Cogn Neurosci Neuroimaging, 3(10), 878–886. 10.1016/j.bpsc.2018.06.007

Smith, S. M., Fox, P. T., Miller, K. L., Glahn, D. C., Fox, P. M., Mackay, C. E., Filippini, N., Watkins, K. E., Toro, R., Laird, A. R., & Beckmann, C. F. (2009). Correspondence of the brain’s functional architecture during activation and rest. Proc Natl Acad Sci U S A, 106(31), 13040–13045. 10.1073/pnas.0905267106

Smith, S. M., Jenkinson, M., Woolrich, M. W., Beckmann, C. F., Behrens, T. E., Johansen-Berg, H., Bannister, P. R., De Luca, M., Drobnjak, I., Flitney, D. E., Niazy, R. K., Saunders, J., Vickers, J., Zhang, Y., De Stefano, N., Brady, J. M., & Matthews, P. M. (2004). Advances in functional and structural MR image analysis and implementation as FSL. Neuroimage, 23 *Suppl 1*, S208–219. 10.1016/j.neuroimage.2004.07.051

Sperling, M. R. (1993). Neuroimaging in epilepsy: recent developments in MR imaging, positron-emission tomography, and single-photon emission tomography [Research Support, U.S. Gov’t, P.H.S. Review]. Neurol Clin, 11(4), 883–903. http://www.ncbi.nlm.nih.gov/pubmed/8272037

Sperling, M. R., Cahan, L. D., & Brown, W. J. (1989). Relief of seizures from a predominantly posterior temporal tumor with anterior temporal lobectomy [Case Reports]. Epilepsia, 30(5), 559–563. 10.1111/j.1528-1157.1989.tb05471.x

Sperling, M. R., O’Connor, M. J., Saykin, A. J., Phillips, C. A., Morrell, M. J., Bridgman, P. A., French, J. A., & Gonatas, N. (1992). A noninvasive protocol for anterior temporal lobectomy [Research Support, U.S. Gov’t, P.H.S.]. Neurology, 42(2), 416–422. 10.1212/wnl.42.2.416

Struck, A. F., Garcia-Ramos, C., Nair, V. A., Prabhakaran, V., Dabbs, K., Boly, M., Conant, L. L., Binder, J. R., Meyerand, M. E., & Hermann, B. P. (2023). The presence, nature and network characteristics of behavioural phenotypes in temporal lobe epilepsy. Brain Commun, 5(2), fcad095. 10.1093/braincomms/fcad095

Szaflarski, J. P., Gloss, D., Binder, J. R., Gaillard, W. D., Golby, A. J., Holland, S. K., Ojemann, J., Spencer, D. C., Swanson, S. J., French, J. A., & Theodore, W. H. (2017). Practice guideline summary: Use of fMRI in the presurgical evaluation of patients with epilepsy: Report of the Guideline Development, Dissemination, and Implementation Subcommittee of the American Academy of Neurology. Neurology, 88(4), 395–402. 10.1212/WNL.0000000000003532

Tai, X. Y., Bernhardt, B., Thom, M., Thompson, P., Baxendale, S., Koepp, M., & Bernasconi, N. (2018). Review: Neurodegenerative processes in temporal lobe epilepsy with hippocampal sclerosis: Clinical, pathological and neuroimaging evidence. Neuropathol Appl Neurobiol, 44(1), 70–90. 10.1111/nan.12458

Tracy, J. I., Chaudhary, K., Modi, S., Crow, A., Kumar, A., Weinstein, D., & Sperling, M. R. (2021). Computational support, not primacy, distinguishes compensatory memory reorganization in epilepsy. Brain Commun, 3(2), fcab025. 10.1093/braincomms/fcab025

Tracy, J. I., Dechant, V., Sperling, M. R., Cho, R., & Glosser, D. (2007). The association of mood with quality of life ratings in epilepsy. Neurology, 68(14), 1101–1107. 10.1212/01.wnl.0000242582.83632.73

Vriend, C., Wagenmakers, M. J., van den Heuvel, O. A., & van der Werf, Y. D. (2020). Resting-state network topology and planning ability in healthy adults. Brain Struct Funct, 225(1), 365–374. 10.1007/s00429-019-02004-6

Wang, D., Buckner, R. L., Fox, M. D., Holt, D. J., Holmes, A. J., Stoecklein, S., Langs, G., Pan, R., Qian, T., Li, K., Baker, J. T., Stufflebeam, S. M., Wang, K., Wang, X., Hong, B., & Liu, H. (2015). Parcellating cortical functional networks in individuals. Nat Neurosci, 18(12), 1853–1860. 10.1038/nn.4164

Wang, Y., & Li, T. Q. (2015). Dimensionality of ICA in resting-state fMRI investigated by feature optimized classification of independent components with SVM. Front Hum Neurosci, 9, 259. 10.3389/fnhum.2015.00259

Ward, J. H. (1963). Hierarchical Grouping to Optimize an Objective Function. Journal of the American Statistical Association, 58(301), 236–244. 10.2307/2282967

Yang, S., Zhang, Z., Chen, H., Meng, Y., Li, J., Li, Z., Xu, Q., Zhang, Q., Fan, Y. S., Lu, G., & Liao, W. (2021). Temporal variability profiling of the default mode across epilepsy subtypes. Epilepsia, 62(1), 61–73. 10.1111/epi.16759

Yeo, B. T., Krienen, F. M., Sepulcre, J., Sabuncu, M. R., Lashkari, D., Hollinshead, M., Roffman, J. L., Smoller, J. W., Zollei, L., Polimeni, J. R., Fischl, B., Liu, H., & Buckner, R. L. (2011). The organization of the human cerebral cortex estimated by intrinsic functional connectivity. J Neurophysiol, 106(3), 1125–1165. 10.1152/jn.00338.2011

Zhang, Z., Liao, W., Xu, Q., Wei, W., Zhou, H. J., Sun, K., Yang, F., Mantini, D., Ji, X., & Lu, G. (2017). Hippocampus-associated causal network of structural covariance measuring structural damage progression in temporal lobe epilepsy. Hum Brain Mapp, 38(2), 753–766. 10.1002/hbm.23415

Zhao, F., Kang, H., You, L., Rastogi, P., Venkatesh, D., & Chandra, M. (2014). Neuropsychological deficits in temporal lobe epilepsy: A comprehensive review. Ann Indian Acad Neurol, 17(4), 374–382. 10.4103/0972-2327.144003

