## Supplementary for "Normative and individual, non-normative intrinsic networks and the transition to impaired cognition"

**Author affiliations:**

Correspondence to: Joseph I. Tracy

### Supplementary Materials and Methods

#### Definition of terms

Cognitively intact patients: cluster 1 in K-means clustering of neuropsychological measures, identifying the group of patient with overall higher performance.

Cognition impaired patients: cluster 2 in K-means cognition clustering of neuropsychological measures, identifying the group of patient with overall lower performance.

Canonical ICNs: Canonical, well-established “normative” intrinsic connectivity networks (ICNs) first reported by Yeo et al. (Yeo et al., 2011) or Smith et al. (Smith et al., 2009)

Individualized ICNs: Individualized ICN component from a participant via independent component analysis.

Normative ICNs: Individualized ICNs that strongly match a given canonical ICN.

Non-normative ICNs: Individualized ICNs that poorly match a given canonical ICN.

Best matching canonical ICNs: Individualized ICNs that best matches each canonical ICN, representing the match degree to each of the canonical ICNs.

Best matching individualized ICNs: Canonical ICNs that best matches each individualized ICN, representing the normativity of a person’s individualized ICNs.

ICN number: the ICN number of normative or non-normative ICNs.

ICN variance explained: Sum of the variance explained of normative or non-normative ICNs.

ICN GM volume: Sum of the gray matter volume underneath ICNs of normative or non-normative ICNs.

ICN strength: Mean intra-ICN connectivity strength across each voxel, summed, in normative or non-normative ICNs.

Network efficiency: Inter-ICN global network efficiency represented the level of functional integration (communication) among the components within ICN type (normative, non-normative).

Vocabulary: Wechsler Adult Intelligence Scale (WAIS) III or IV(Franz Petermann, 2008) vocabulary subtest.

Similarities: WAIS-III or IV Similarities subtest.

Semantic fluency: Controlled oral word association test(Benton et al., 1994; Gladsjo et al., 1999) -- semantic fluency.

Letter fluency: Controlled oral word association test(Benton et al., 1994; Gladsjo et al., 1999) -- letter fluency.

BNT: Boston naming test(Goodglass et al., 2001).

Matrix Reasoning: WAIS-III or IV(Franz Petermann, 2008) matrix reasoning.

Digit Span: WAIS-III or IV(Franz Petermann, 2008) Digit Span subtest

WCST PR: Wisconsin card sort test(Barcelo et al., 1997) -- perseverative responses

WCST CC: Wisconsin card sort test(Barcelo et al., 1997) -- categories completed

TMTB: Trail making test(Bowie & Harvey, 2006) -- part B

TMTA: Trail making test(Bowie & Harvey, 2006) -- part A

Coding: WAIS-III or IV(Franz Petermann, 2008) digit symbol coding subtest

Logical Memory 1: Wechsler memory sale III or IV(WECHSLER, 2022) logical memory 1

Logical Memory 2: Wechsler memory scale III or IV(WECHSLER, 2022) logical memory 2

CVLT TL: California verbal learning test(Delis, 2000) -- total learning

CVLT TLDFR: California verbal learning test(Delis, 2000) -- Long Delay Free Recall

Block Design: WAIS-III or IV(Franz Petermann, 2008) Block Design subtest

ROCF Copy: Rey–Osterrieth complex figure test(Shin et al., 2006) -- copy condition

Pegboard R: Grooved pegboard(Skogan et al., 2018) -- right hand

Pegboard L: Grooved pegboard(Skogan et al., 2018) -- left hand

#### Imaging Methods

All participants were scanned on a 3-T X-series Philips Achieva clinical MRI scanner (Amsterdam, the Netherlands) using an 8-channel head coil. A 5-minute functional MRI scan involving a resting-state condition was collected from all participants. During the resting-state condition, participants viewed a crosshair with no task requirements and were instructed to remain still throughout the scan and not fall asleep. After the scan, all participants reported that they were able to complete the scan as instructed. The fMRI data was collected with a single shot echoplanar gradient echo imaging (EPI) sequence acquiring T2* signals (120 - 480 volumes; 34 axial slices acquired parallel to the anterior, posterior commissure line; TR = 2.5 s, TE = 35 ms; FOV = 256 mm, 128 × 128 data matrix voxels, flip angle = 90°, in-plane resolution = 2 mm × 2 mm, slice thickness = 4 mm). Each EPI imaging series started with three discarded scans to allow for signal stabilization Prior to collection of the T2* images, T1-weighted images (180 slices) were collected using an MPRAGE sequence (256 × 256 isotropic 1mm voxels; TR = 640 ms; TE = 3.2 ms, FOV = 256 mm, flip angle = 8°) in positions identical to the functional scans to provide an anatomical reference. The in-plane resolution for each T1-weighted slice was 1 mm3 (axial oblique). The raw dicoms were converted to nifti format using the dcm2niix software tool. The nifti files were then renamed in BIDS standard for batch level pre-processing.

#### Image data processing

FMRI and T1w-weighted data were preprocessed using *fMRIPrep* (version 22.1.1, [RRID: SCR_016216] (Esteban et al., 2019), based on *Nipype* 1.8.5 [RRID:SCR_002502])(Gorgolewski et al., 2011).

##### Anatomical data preprocessing

The T1-weighted (T1w) image was corrected for intensity non-uniformity (INU) with N4BiasFieldCorrection (Tustison et al., 2010), distributed with ANTs 2.3.3, and used as T1w-reference throughout the workflow. The T1w-reference was then skull-stripped with a Nipype implementation of the antsBrainExtraction.sh workflow (from ANTs), using OASIS30ANTs as target template. Brain tissue segmentation of cerebrospinal fluid (CSF), white-matter (WM) and gray-matter (GM) was performed on the brain-extracted T1w using fast (FSL 6.0.5) (Zhang et al., 2001). Brain surfaces were reconstructed using recon-all (FreeSurfer 7.2.0, RRID:SCR_001847, Dale, Fischl, and Sereno, 1999), and the brain mask estimated previously was refined with a custom variation of the method to reconcile ANTs-derived and FreeSurfer-derived segmentations of the cortical gray-matter of Mindboggle (RRID:SCR_002438, Klein et al. 2017).

##### Functional data preprocessing

For each participant, the following preprocessing steps were performed. A deformation field to correct for susceptibility distortions was estimated based on fMRIPrep’s fieldmap-less approach. The deformation field resulted from co-registering the EPI reference to the same-subject’s T1w-reference with its intensity inverted (Wang et al., 2017). Registration was performed with antsRegistration (ANTs 2.3.3), and the process regularized by constraining deformation to be nonzero only along the phase-encoding direction, and modulated with an average fieldmap template (Treiber et al., 2016). A reference volume (a median of motion corrected subset of volumes) and its skull-stripped version were generated using a custom methodology of *fMRIPrep*. Head-motion parameters with respect to the BOLD reference image (transformation matrices, and six corresponding rotation and translation parameters) were estimated before any spatiotemporal filtering using mcflirt (FSL 6.0.5). BOLD runs were slice-time corrected to 1.22s (0.5 of slice acquisition range 0s-2.43s) using 3dTshift from AFNI (RRID:SCR_005927) (Cox & Hyde, 1997) . The BOLD time-series (including slice-timing correction when applied) were resampled onto their original, native space by applying the transforms to correct for head-motion. These resampled BOLD time-series will be referred to as *preprocessed BOLD in original space*, or just *preprocessed BOLD*.

The BOLD reference was then co-registered to the T1w reference using bbregister (FreeSurfer) which implements boundary-based registration(Dale et al., 1999).

Co-registration was configured with six degrees of freedom. Several confounding time-series were calculated based on the *preprocessed BOLD*: framewise displacement (FD), DVARS and three region-wise global signals. FD was computed using two formulations following Power (absolute sum of relative motions) (Power et al., 2014) and Jenkinson (relative root mean square displacement between affines) (Jenkinson et al., 2002). FD and DVARS are calculated for each functional run, both using their implementations in *Nipype*. The three global signals are extracted within the CSF, the WM, and the whole-brain masks. The head-motion estimates calculated in the correction step were placed within the corresponding confounds file. The confound time series derived from head motion estimates and global signals were expanded with the inclusion of temporal derivatives and quadratic terms for each (Satterthwaite et al., 2013a). Automatic removal of motion artifacts using independent component analysis (ICA-AROMA(Pruim et al., 2015)) was performed on the preprocessed BOLD on MNI space time-series after removal of non-steady state volumes and spatial smoothing with an isotropic, Gaussian kernel of 6mm FWHM (full-width half-maximum). Corresponding “non-aggressively” denoised runs were produced after such smoothing. Additionally, the “aggressive” noise-regressors were collected and placed in the corresponding confounds file.

After *fMRIPrep* estimation of head-motion, we excluded 13 TLE patient with minimum continuous 120 frame average head movement larger than 0.4. Thus, 63 cognitively intact and 24 cognitively impaired patients will included in following image analysis.

The eXtensible Connectivity Pipeline (XCP) (Ciric et al., 2018; Satterthwaite et al., 2013b) was used to post-process the outputs of fMRIPrep version 22.1.1. XCP was built with Nipype 1.8.5. The native space BOLD fMRI data and confounds were mean-centered and linearly detrended. AROMA motion-labeled components(Pruim et al., 2015), mean white matter signal, and mean CSF signal were selected as nuisance regressors(Ciric et al., 2018; Satterthwaite et al., 2013b). AROMA non-motion components (i.e., ones assumed to reflect signal) were also included in the regression, as implemented in nilearn 0.9.2(Abraham et al., 2014), after which the resulting parameter estimates from only the nuisance regressors were used to denoise the BOLD data. In this way, shared variance between the nuisance regressors and the signal regressors was separated smartly, so that signal would not be removed by the regression. This is colloquially known as ‘non-aggressive’ denoising, and is the recommended denoising method when nuisance regressors may share variance with known signal regressors(Pruim et al., 2015). The interpolated time series were then band-pass filtered to retain signals within the 0.009-0.08 Hz frequency band. The processed BOLD was smoothed using Nilearn with a gaussian kernel size of 6.0 mm (FWHM) and resampled into standard space, generating a preprocessed BOLD run in 2mm MNI space.

**ICN measures for the normative and non-normative ICNs**

ICN GM volume measure we calculated the GM volume underlying the regions in each ICN then summed the volume within each ICN type(normative, non-normative). Prior to this, each component was thresholded (FSL default level of 0.5 or greater), and the resulting mask, containing the ICN regions, was then overlaid on the person-specific gray matter probability segmentation map. The participant’s individual total brain GM volume was used to normalize each of the ICN GM volume calculations, allowing us to account for individual differences in brain size.

ICN strength represented the strength of functional connectivity (edges) within each component, reflecting the level of intrinsic communication among the regions in a given ICN component. To compute this, we first using the same thresholded ICN mask noted above, calculated the functional connectivity (Pearson correlation) between all the voxels underling a given ICN component mask. Next, we calculated the mean of the inter-regional voxel-wise correlations for each ICN component for each participant. The total ICN strength is the sum of the ICN strength within the ICN type (normative, non-normative).

The overall efficiency between the components within each ICN type represented the level of functional integration (communication) among the components within ICN type (normative, non-normative). Efficiency was defined as inversely proportion to the harmonic mean of the shortest distance (i.e. number of edges) between all possible pairs of nodes/networks (Latora & Marchiori, 2001) .We first calculated the functional connectivity between ICNs by Pearson correlation. Using the Brain Connectivity Toolbox(Rubinov & Sporns, 2010) (<https://sites.google.com/site/bctnet/home>), efficiency was then calculated using a binarized connection matrix containing the top 10% of edge connections (Achard & Bullmore, 2007; Kim et al., 2017).

### Supplementary Results

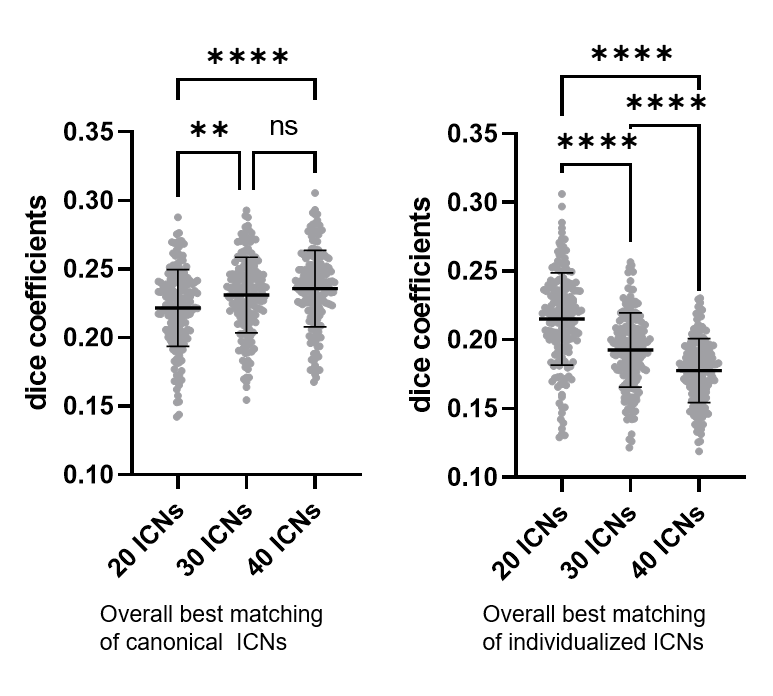

**Supplementary Figure 1. Matching level cross different individualized ICN number.** Each point represents the average best matching value for canonical ICNs or individualized ICNs for each subjects, using the Yeo networks as an example. It can be observed that as the number of individualized ICNs increases average best matching of canonical ICNs gradually increases. However, there was no significant improvement after 30 ICN. At the same time the average best matching of individualized ICN gradually decreases. This is because the increase in the number of individualized ICNs leads to the isolation of more non-normative components. Thirty individualized ICNs compared to the other choices largely balances the mutual matching between individualized ICN and normative ICNs. ns: *p* > 0.05, *: *p* < 0.05, **: *p* < 0.01, ***: *p* < 0.001, ****: *p* < 0.0001.

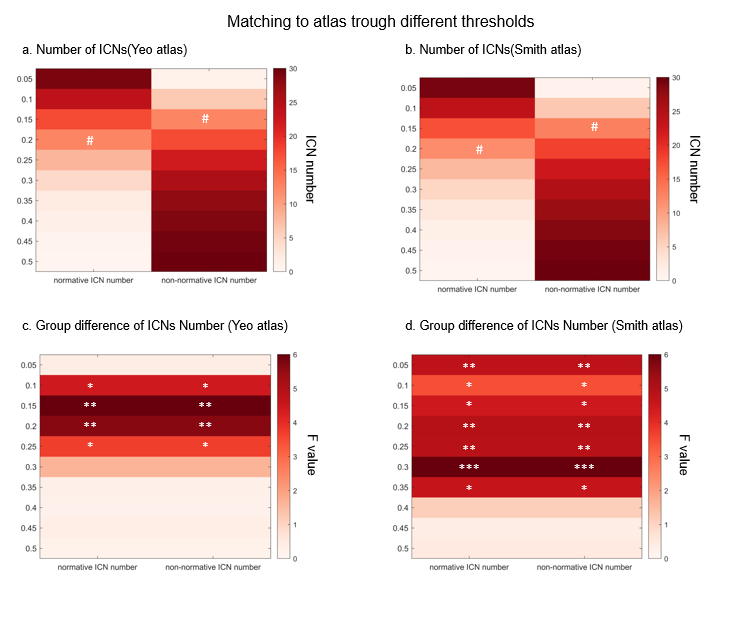

**Supplementary Figure 2.** **Matching to atlas trough different thresholds.** Normative ICN number and non-normative ICN number changes with threshold in Yeo atlas **(a)** and in Smith atlas **(b)**. Optimal normative ICN threshold was 0.2, while the non-normative ICN threshold was 0.15 for both atlases, marked as #. The two classes of ICNs showed roughly equal numbers at the optimal thresholds. We also validated the stability of the between-group differences under different matching thresholds using number of ICNs (one-way ANOVA). The number of ICNs showed stable between-group differences at dice coefficients of 0.1-0.25 for the Yeo atlas **(c)**, and spatial correlation coefficients of 0.05-0.35 for the Smith atlas **(d)**. *: *p* < 0.05, **: *p* < 0.01, ***: *p* < 0.001.

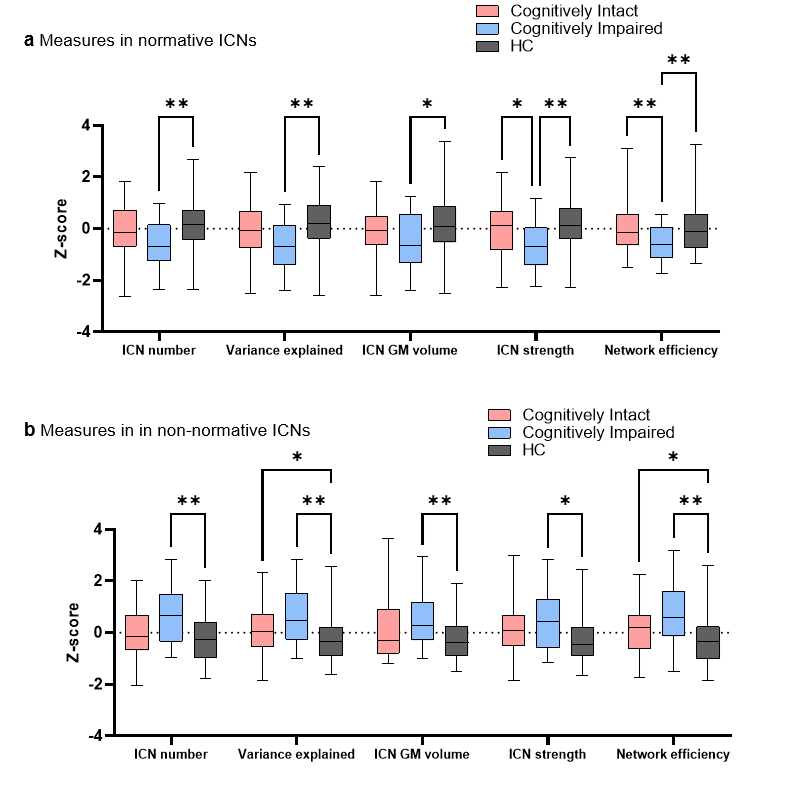

**Supplementary Figure 3. Two-way mixed measures ANOVA on each ICN type (Figure 3bc).** Box plots are used here for more complete data distribution and statistical significance for ICN measures of normative ICNs **(a)**, and non-normative ICNs **(b)**. *: *p* < 0.05, **: *p* < 0.01, Tukey’s corrected.

**Supplementary Table 1. Two-way mixed measures ANOVA on matching individualized ICNs to canonical ICNs (Figure 3a)**

| **Two-way mixed measures ANOVA** | **Sums of Squares** | **Degree of Freedom** | ***F* / Tukey's *Q*** | ***p* / Adjusted *p*** |
| --- | --- | --- | --- | --- |
| **Main effects** |  |  |  |  |
| Network x Group | 0.1967 | 32 | 1.357 | 0.0874 |
| Network | 14.12 | 16 | 194.8 | 0.0001 |
| Group | 0.1573 | 2 | 6.445 | 0.0020 |
| Subject | 2.148 | 176 | 2.693 | 0.0001 |
| Residual | 12.76 | 2816 |  |  |
| **Post-hoc test of main group effects** |  |  |  |  |
| Impaired vs. Intact |  |  | 3.644 | 0.0289 |
| Intact vs. HC |  |  | 1.759 | 0.4292 |
| Impaired vs. HC |  |  | 5.068 | 0.0013 |
| **Post-hoc test of group effect in each network** |  |  |  |  |
| visualCentral |  |  |  |  |
| Impaired vs. Intact |  |  | 0.9167 | 0.7946 |
| Intact vs. HC |  |  | 2.02 | 0.3294 |
| Impaired vs. HC |  |  | 2.167 | 0.2904 |
| visualPeripheral |  |  |  |  |
| Impaired vs. Intact |  |  | 2.29 | 0.2502 |
| Intact vs. HC |  |  | 0.8448 | 0.8218 |
| Impaired vs. HC |  |  | 2.926 | 0.1125 |
| somatomotorA |  |  |  |  |
| Impaired vs. Intact |  |  | 2.273 | 0.2531 |
| Intact vs. HC |  |  | 1.933 | 0.3615 |
| Impaired vs. HC |  |  | 0.9553 | 0.7791 |
| Impaired vs. Intact |  |  | 1.8 | 0.4186 |
| Intact vs. HC |  |  | 1.55 | 0.5183 |
| Impaired vs. HC |  |  | 0.8567 | 0.818 |
| dorsalAttentionA |  |  |  |  |
| Impaired vs. Intact |  |  | 5.384 | 0.0012 |
| Intact vs. HC |  |  | 1.844 | 0.3957 |
| Impaired vs. HC |  |  | 4.28 | 0.012 |
| dorsalAttentionB |  |  |  |  |
| Impaired vs. Intact |  |  | 1.535 | 0.5285 |
| Intact vs. HC |  |  | 0.4674 | 0.9416 |
| Impaired vs. HC |  |  | 1.911 | 0.3764 |
| Impaired vs. Intact |  |  | 1.313 | 0.626 |
| Intact vs. HC |  |  | 2.866 | 0.1098 |
| Impaired vs. HC |  |  | 0.5215 | 0.928 |
| Impaired vs. Intact |  |  | 0.2332 | 0.9851 |
| Intact vs. HC |  |  | 0.07903 | 0.9983 |
| Impaired vs. HC |  |  | 0.1889 | 0.9902 |
| limbicOrbitofrontal |  |  |  |  |
| Impaired vs. Intact |  |  | 0.12 | 0.996 |
| Intact vs. HC |  |  | 0.1572 | 0.9932 |
| Impaired vs. HC |  |  | 0.02527 | 0.9998 |
| limbicTemporopolar |  |  |  |  |
| Impaired vs. Intact |  |  | 5.88 | 0.0002 |
| Intact vs. HC |  |  | 0.0657 | 0.9988 |
| Impaired vs. HC |  |  | 6.588 | <0.0001 |
| frontoparietalControlA |  |  |  |  |
| Impaired vs. Intact |  |  | 0.4599 | 0.9435 |
| Intact vs. HC |  |  | 1.779 | 0.4219 |
| Impaired vs. HC |  |  | 0.7152 | 0.8691 |
| frontoparietalControlB |  |  |  |  |
| Impaired vs. Intact |  |  | 0.5918 | 0.9082 |
| Intact vs. HC |  |  | 1.9 | 0.3736 |
| Impaired vs. HC |  |  | 1.714 | 0.4553 |
| frontoparietalControlC |  |  |  |  |
| Impaired vs. Intact |  |  | 1.227 | 0.6632 |
| Intact vs. HC |  |  | 2.572 | 0.1672 |
| Impaired vs. HC |  |  | 3.222 | 0.0704 |
| defaultA |  |  |  |  |
| Impaired vs. Intact |  |  | 1.312 | 0.6264 |
| Intact vs. HC |  |  | 1.97 | 0.3474 |
| Impaired vs. HC |  |  | 2.71 | 0.1487 |
| defaultB |  |  |  |  |
| Impaired vs. Intact |  |  | 0.02475 | 0.9998 |
| Intact vs. HC |  |  | 2.282 | 0.2432 |
| Impaired vs. HC |  |  | 1.675 | 0.4698 |
| defaultC |  |  |  |  |
| Impaired vs. Intact |  |  | 1.824 | 0.4102 |
| Intact vs. HC |  |  | 0.7274 | 0.8645 |
| Impaired vs. HC |  |  | 2.369 | 0.2327 |
| temporoparietal |  |  |  |  |
| Impaired vs. Intact |  |  | 0.6656 | 0.8853 |
| Intact vs. HC |  |  | 1.964 | 0.35 |
| Impaired vs. HC |  |  | 2.356 | 0.2313 |

**Supplementary Table 2. Two-way mixed measures ANOVA on ICN measures of normative ICNs (Figure 3b)**

| **Two-way mixed measures ANOVA** | **Sums of Squares** | **Degree of Freedom** | ***F* / Tukey's *Q*** | ***p* / Adjusted *p*** |
| --- | --- | --- | --- | --- |
| **Main effects** |  |  |  |  |
| Measure x Group | 3.763 | 8 | 1.332 | 0.2241 |
| Measure | 0.2853 | 4 | 0.2020 | 0.7876 |
| Group | 55.77 | 2 | 8.434 | 0.0003 |
| Subject | 581.8 | 176 | 9.361 | 0.0001 |
| Residual | 248.6 | 704 |  |  |
| **Post-hoc test of main group effects** |  |  |  |  |
| Impaired vs. Intact |  |  | 3.899 | 0.0176 |
| Intact vs. HC |  |  | 2.359 | 0.2203 |
| Impaired vs. HC |  |  | 5.764 | 0.0002 |
| **Post-hoc test of group effect in each measure** |  |  |  |  |
| ICN number |  |  |  |  |
| Impaired vs. Intact |  |  | 3.05 | 0.0909 |
| Intact vs. HC |  |  | 3.036 | 0.0846 |
| Impaired vs. HC |  |  | 5.356 | 0.0016 |
| Variance explained |  |  |  |  |
| Impaired vs. Intact |  |  | 3.276 | 0.0644 |
| Intact vs. HC |  |  | 2.665 | 0.1473 |
| Impaired vs. HC |  |  | 5.35 | 0.0017 |
| ICN GM volume |  |  |  |  |
| Impaired vs. Intact |  |  | 2.105 | 0.3082 |
| Intact vs. HC |  |  | 2.419 | 0.2051 |
| Impaired vs. HC |  |  | 3.706 | 0.0343 |
| ICN strength | |  |  |  |
| Impaired vs. Intact |  |  | 3.81 | 0.0265 |
| Intact vs. HC |  |  | 1.985 | 0.3422 |
| Impaired vs. HC |  |  | 5.525 | 0.0012 |
| Network Efficiency |  |  |  |  |
| Impaired vs. Intact |  |  | 4.416 | 0.0078 |
| Intact vs. HC |  |  | 0.1839 | 0.9907 |
| Impaired vs. HC |  |  | 4.575 | 0.0062 |

**Supplementary Table 3. Two-way mixed measures ANOVA on ICN measures of non-normative ICNs (Figure 3c)**

| **Two-way mixed measures ANOVA** | **Sums of Squares** | **Degree of Freedom** | ***F* / Tukey's *Q*** | ***p* / Adjusted *p*** |
| --- | --- | --- | --- | --- |
| **Main effects** |  |  |  |  |
| Measure x Group | 1.711 | 8 | 0.6061 | 0.7732 |
| Measure | 0.4253 | 4 | 0.3012 | 0.7678 |
| Group | 76.82 | 2 | 12.01 | 0.0001 |
| Subject | 563 | 176 | 9.063 | 0.0001 |
| Residual | 248.5 | 704 |  |  |
| **Post-hoc test of main group effects** |  |  |  |  |
| Impaired vs. Intact |  |  | 3.568 | 0.0334 |
| Intact vs. HC |  |  | 3.99 | 0.0147 |
| Impaired vs. HC |  |  | 6.58 | <0.0001 |
| **Post-hoc test of group effect in each measure** |  |  |  |  |
| ICN number |  |  |  |  |
| Impaired vs. Intact |  |  | 3.279 | 0.0651 |
| Intact vs. HC |  |  | 3.267 | 0.0581 |
| Impaired vs. HC |  |  | 5.587 | 0.0011 |
| Variance explained |  |  |  |  |
| Impaired vs. Intact |  |  | 2.863 | 0.1207 |
| Intact vs. HC |  |  | 3.477 | 0.0404 |
| Impaired vs. HC |  |  | 5.117 | 0.003 |
| ICN GM volume |  |  |  |  |
| Impaired vs. Intact |  |  | 2.087 | 0.3113 |
| Intact vs. HC |  |  | 2.703 | 0.1404 |
| Impaired vs. HC |  |  | 4.484 | 0.009 |
| ICN strength | |  |  |  |
| Impaired vs. Intact |  |  | 1.944 | 0.365 |
| Intact vs. HC |  |  | 3.234 | 0.0611 |
| Impaired vs. HC |  |  | 3.93 | 0.0241 |
| Network Efficiency |  |  |  |  |
| Impaired vs. Intact |  |  | 3.058 | 0.0925 |
| Intact vs. HC |  |  | 4.154 | 0.0108 |
| Impaired vs. HC |  |  | 5.551 | 0.0013 |

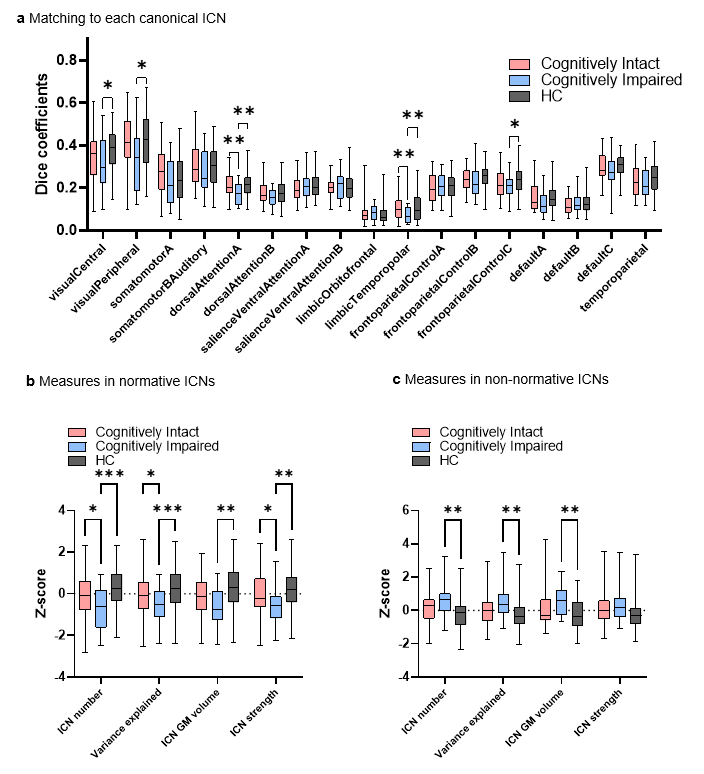

**Supplementary Figure 4. Matching 20 individualized ICNs with canonical ICNs in Yeo 17 atlas.** To show the stability of the main results in terms of the number of individual ICNs, we mixed the main results in Figure 3 using 20 individual ICNs. **(a)** Shows significant decrease in dice coefficients in cognitively impaired patients by two-way mixed measures ANOVA (Group effect: *F* = 8.346 *p* = 0.0003, Impaired vs. Intact *p* = 0.0073, intact vs. HC *p* = 0.4280, and impaired vs. HC *p* = 0.0002, Tukey’s corrected), especially in limbic network temporopolar part, dorsal attention network part A, central visual, peripheral visual, and frontoparietal control part C. **(b)** Box plots shows all the post-hoc analysis statistics in two-way mixed measures ANOVA on normative ICN measures. In general, measures of normative ICNs significantly decreased in cognitively impaired patients (Group effect: *F* = 9.417 *p* = 0.0001, Impaired vs. Intact *p* = 0.0199, intact vs. HC *p* = 0.1155 and impaired vs. HC *p* < 0.0001, Tukey’s corrected). **(c)** Same box plots but in non-normative ICNs. In general, measures of non-normative ICNs significantly increased in cognitively impaired patients compared to controls (Group effect: *F* = 7.752, *p* = 0.0006, Impaired vs. Intact *p* = 0.1299, intact vs. HC *p* = 0.0522 and impaired vs. HC *p* = 0.008, Tukey’s corrected). Because some patients did not have enough non-normative ICNs to calculate network efficiency, we did not include this measure in this validation. *: *p* < 0.05, **: *p* < 0.01, ***: *p* < 0.001, Tukey’s corrected.

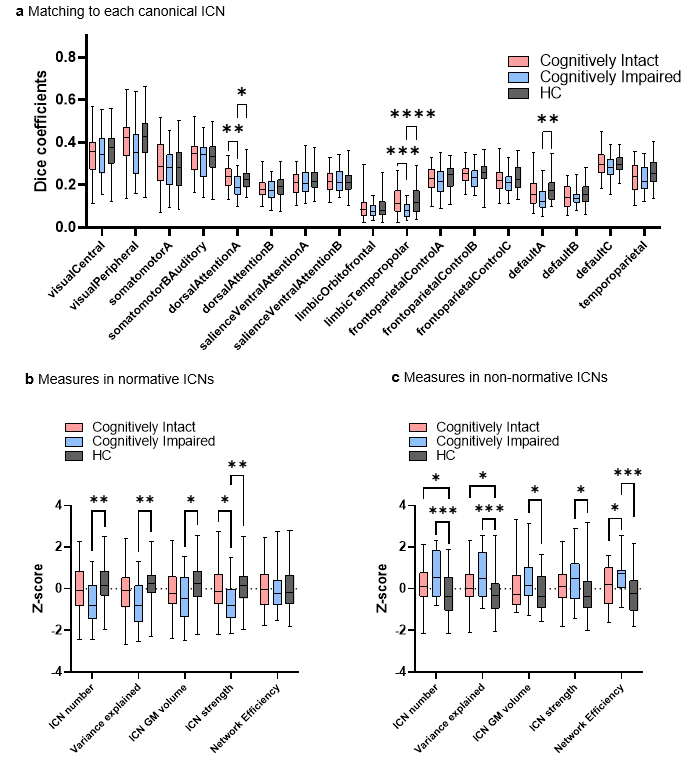

**Supplementary Figure 5. Matching 40 individualized ICNs with canonical ICNs in Yeo 17 atlas.** To show the stability of the main results in terms of the number of individual ICNs, we mixed the main results in Figure 3 using 40 individual ICNs. **(a)** Shows significantly decreased dice coefficients in cognitively impaired patients by two-way mixed measures ANOVA (Group effect: *F* = 7.027 *p* = 0.0012, Impaired vs. Intact *p* = 0.0276, intact vs. HC *p* = 0.3305 and impaired vs. HC *p* = 0.0007, Tukey’s corrected), especially in limbic network temporopolar part, dorsal attention network part A, and default part A. **(b)** Box plots shows all the post-hoc analysis statistics in two-way mixed measures ANOVA on measures of normative ICNs. In general, measures of normative ICNs significantly decreased in cognitively impaired patients (Group effect: *F* = 6.292 *p* = 0.0023, Impaired vs. Intact *p* = 0.0446, intact vs. HC p = 0.3399 and impaired vs. HC *p* = 0.0016, Tukey’s corrected). **(c)** Same box plots but in non-normative ICNs. In general, measures of non-normative ICNs significantly increased in cognitively impaired patients compare to controls (Group effect: *F* = 12.30, *p* < 0.0001, Impaired vs. Intact *p* = 0.0463, intact vs. HC *p* = 0.0086 and impaired vs. HC *p* < 0.0001, Tukey’s corrected). *: *p* < 0.05, **: *p* < 0.01, ***: *p* < 0.001, ****: *p* < 0.0001, Tukey’s corrected.

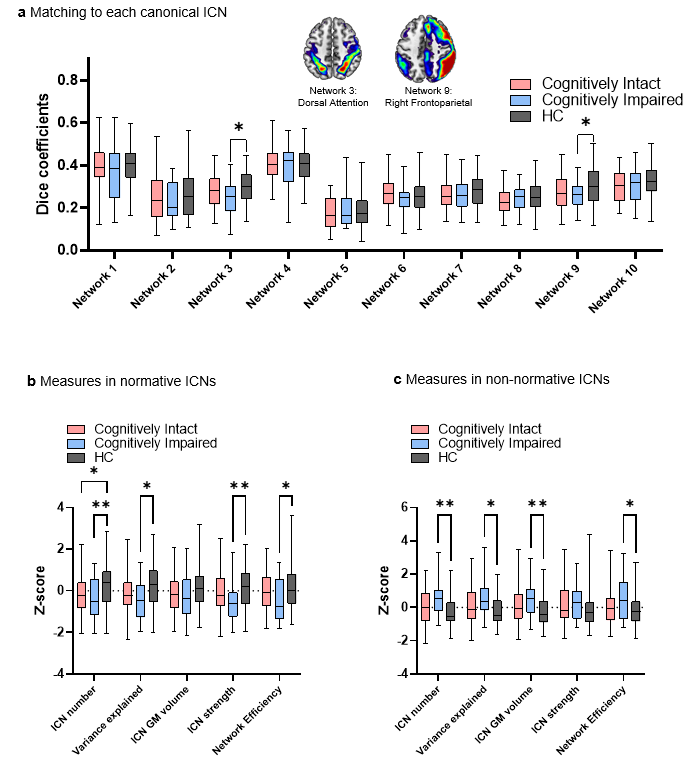

**Supplementary Figure 6. Matching 30 individualized ICNs with canonical ICNs in Smith 10 networks atlas.** To show the stability of the main results in terms of the different atlas and match strategy, we mixed the main results in Figure 3 using measures yielded by match to Smith 10 networks. **(a)** Shows significantly decreased dice coefficients in cognitively impaired patients compare to controls by two-way mixed measures ANOVA (Group effect: *F* = 5.464 *p* = 0.0050, Impaired vs. Intact *p* = 0.5053, intact vs. HC *p* = 0.0464 and impaired vs. HC *p* = 0.0125, Tukey’s corrected), especially in dorsal attention network, and right frontoparietal network. **(b)** Box plots shows all the post-hoc analysis statistics in two-way mixed measures ANOVA on measures of normative ICNs. In general, measures of normative ICNs significantly decreased in cognitively impaired patients (Group effect: *F* = 6.370 *p* = 0.0021, Impaired vs. Intact *p* = 0.2606, intact vs. HC *p* = 0.0592 and impaired vs. HC *p* = 0.0035, Tukey’s corrected). **(c)** Same box plots but in non-normative ICNs. In general, measures of non-normative ICNs significantly increased in cognitively impaired patients compare to controls (Group effect: *F* = 7.561, *p* = 0.0007, Impaired vs. Intact *p* = 0.0865, intact vs. HC *p* = 0.0903 and impaired vs. HC *p* = 0.0007, Tukey’s corrected). *: *p* < 0.05, **: *p* < 0.01, Tukey’s corrected.

**Supplementary Table 4. PLS-SEM analysis of ICN measure influence on cognitive performance (Figure 4a)**

|  | **Outer Loading** | **bootstrapping mean** | **bootstrapping deviation** | ***t* statistics** | ***p* values** |
| --- | --- | --- | --- | --- | --- |
| **ICN measures** |  |  |  |  |  |
| Normative ICNs number | 0.919 | 0.874 | 0.181 | 5.071 | <0.001 |
| Non-normative ICNs number | -0.898 | -0.852 | 0.182 | 4.948 | <0.001 |
| Normative ICNs variance explained | 0.914 | 0.87 | 0.176 | 5.189 | <0.001 |
| Non-normative ICNs variance explained | -0.895 | -0.847 | 0.182 | 4.903 | <0.001 |
| Normative ICNs GM volume | 0.703 | 0.667 | 0.177 | 3.981 | <0.001 |
| Non-normative ICNs GM volume | -0.543 | -0.518 | 0.132 | 4.126 | <0.001 |
| Normative ICNs strength | 0.923 | 0.882 | 0.164 | 5.62 | <0.001 |
| Non-normative ICNs strength | -0.747 | -0.703 | 0.173 | 4.323 | <0.001 |
| Normative ICNs network efficiency | 0.394 | 0.385 | 0.129 | 3.052 | 0.002 |
| Non-normative ICNs network efficiency | -0.643 | -0.611 | 0.164 | 3.933 | <0.001 |
| Best matching canonical ICNs | 0.919 | 0.874 | 0.169 | 5.433 | <0.002 |
| Best matching individualized ICNs | 0.958 | 0.912 | 0.172 | 5.563 | <0.003 |
| Match and Variance coupling | 0.564 | 0.543 | 0.108 | 5.214 | <0.004 |
| VisualCentral | 0.55 | 0.531 | 0.123 | 4.474 | <0.005 |
| VisualPeripheral | 0.576 | 0.558 | 0.129 | 4.483 | <0.006 |
| SomatomotorA | 0.531 | 0.508 | 0.125 | 4.246 | <0.007 |
| SomatomotorBAuditory | 0.448 | 0.42 | 0.162 | 2.764 | 0.006 |
| DorsalAttentionA | 0.603 | 0.589 | 0.125 | 4.822 | <0.001 |
| DorsalAttentionB | 0.515 | 0.482 | 0.13 | 3.956 | <0.001 |
| SalienceVentralAttentionA | 0.264 | 0.236 | 0.178 | 1.481 | 0.139 |
| SalienceVentralAttentionB | 0.188 | 0.172 | 0.168 | 1.119 | 0.263 |
| LimbicOrbitofrontal | 0.116 | 0.106 | 0.141 | 0.823 | 0.41 |
| LimbicTemporopolar | 0.158 | 0.156 | 0.12 | 1.317 | 0.188 |
| FrontoparietalControlA | 0.108 | 0.098 | 0.133 | 0.81 | 0.418 |
| FrontoparietalControlB | 0.395 | 0.369 | 0.182 | 2.169 | 0.03 |
| FrontoparietalControlC | 0.255 | 0.239 | 0.156 | 1.634 | 0.102 |
| DefaultA | 0.517 | 0.494 | 0.134 | 3.865 | <0.001 |
| DefaultB | -0.001 | -0.006 | 0.144 | 0.004 | 0.997 |
| DefaultC | 0.247 | 0.241 | 0.119 | 2.068 | 0.039 |
| Temporoparietal | 0.407 | 0.38 | 0.157 | 2.593 | 0.01 |
| **neurocognitive performance** |  |  |  |  |  |
| Vocabulary | 0.653 | 0.629 | 0.159 | 4.114 | <0.001 |
| Similarities | 0.697 | 0.672 | 0.132 | 5.277 | <0.001 |
| Semantic fluency | 0.578 | 0.554 | 0.14 | 4.132 | <0.001 |
| Letter Fluency | 0.725 | 0.697 | 0.118 | 6.121 | <0.001 |
| BNT | 0.606 | 0.581 | 0.131 | 4.633 | <0.001 |
| Matrix Reasoning | 0.498 | 0.477 | 0.158 | 3.161 | 0.002 |
| Digit Span | 0.692 | 0.665 | 0.126 | 5.474 | <0.001 |
| WCST PR | 0.288 | 0.281 | 0.147 | 1.961 | 0.05 |
| WCST CC | 0.277 | 0.266 | 0.167 | 1.655 | 0.098 |
| TMTB | 0.648 | 0.617 | 0.128 | 5.069 | <0.001 |
| TMTA | 0.554 | 0.534 | 0.137 | 4.051 | <0.002 |
| Coding | 0.596 | 0.562 | 0.13 | 4.597 | <0.003 |
| Logical Memory 1 | 0.571 | 0.547 | 0.162 | 3.527 | <0.004 |
| Logical Memory 2 | 0.588 | 0.566 | 0.155 | 3.788 | <0.005 |
| CVLT TL | 0.69 | 0.656 | 0.155 | 4.468 | <0.006 |
| CVLT TLDFR | 0.714 | 0.686 | 0.153 | 4.658 | <0.007 |
| Block Design | 0.493 | 0.465 | 0.162 | 3.044 | 0.002 |
| ROCF Copy | 0.418 | 0.4 | 0.168 | 2.483 | 0.013 |
| Pegboard R | 0.502 | 0.473 | 0.173 | 2.909 | 0.004 |
| Pegboard L | 0.43 | 0.404 | 0.17 | 2.538 | 0.011 |

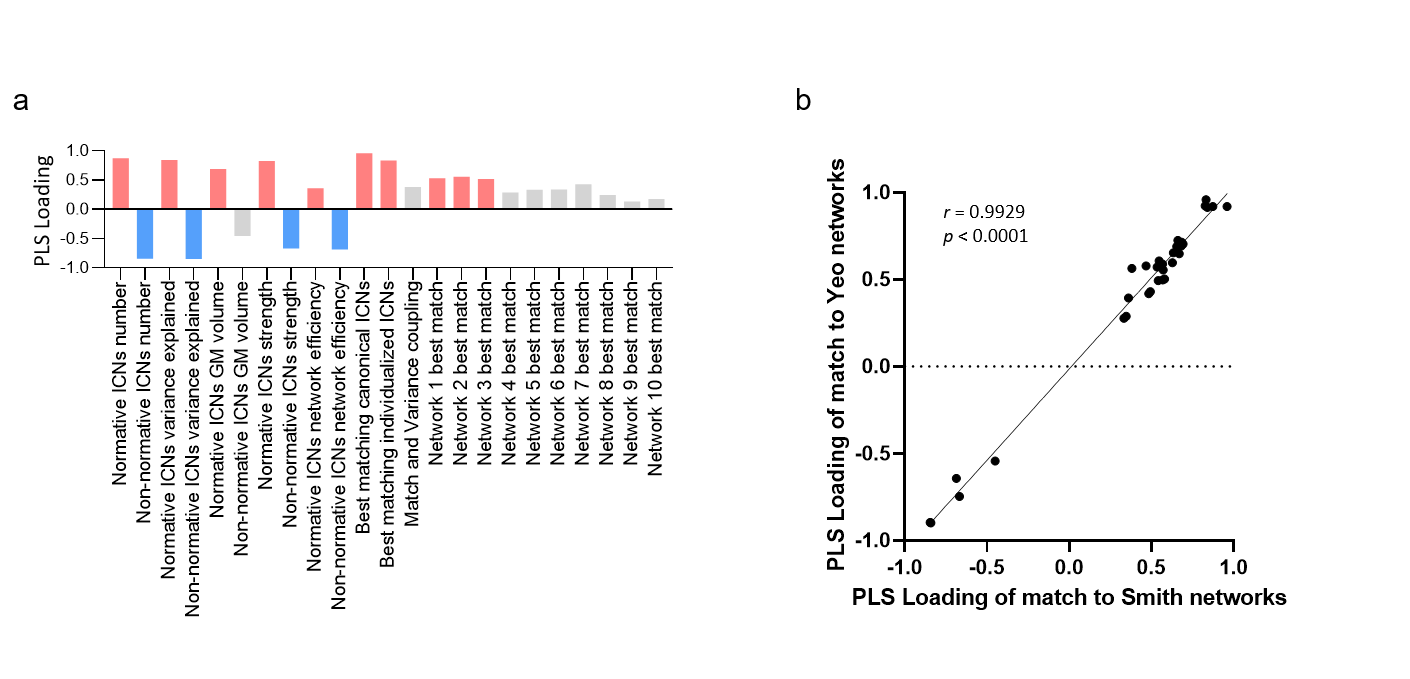

**Supplementary Figure 7. Relationship to neurocognitive performance using Smith 10 networks atlas.** To show the stability of relationship between match measures and cognitive performance, we repeated these results using measures yielded by match to Smith 10 networks. **(a)** The y-axis of the histogram is the PLS-SEM loading of the ICN measures in the model. Red represents measures with significant positive loading, blue represents measures with significant negative loading, and gray represents measures have no significant loading. **(b)** The loadings of match to Yeo networks are highly correlated with the loadings of match to Smith networks.

**Supplementary Table 5. Comparison of outer Loadings of Yeo and Smith PLS-SEM models.**

|  | **Outer Loadings of Yeo network analysis** | **Outer Loadings of Smith network analysis** | **Difference** |
| --- | --- | --- | --- |
| Normative ICNs number | 0.919 | 0.873 | 0.046 |
| Non-normative ICNs number | -0.898 | -0.841 | -0.057 |
| Normative ICNs variance explained | 0.914 | 0.84 | 0.074 |
| Non-normative ICNs variance explained | -0.895 | -0.845 | -0.05 |
| Normative ICNs GM volume | 0.703 | 0.692 | 0.011 |
| Non-normative ICNs GM volume | -0.543 | -0.451 | -0.092 |
| Normative ICNs strength | 0.923 | 0.826 | 0.097 |
| Non-normative ICNs strength | -0.747 | -0.668 | -0.079 |
| Normative ICNs network efficiency | 0.394 | 0.36 | 0.034 |
| Non-normative ICNs network efficiency | -0.643 | -0.687 | 0.044 |
| Best matching canonical ICNs | 0.919 | 0.96 | -0.041 |
| Best matching individualized ICNs | 0.958 | 0.832 | 0.126 |
| Match and Variance coupling | 0.564 | 0.381 | 0.183 |
| Letter Fluency | 0.725 | 0.66 | 0.065 |
| CVLT TLDFR | 0.714 | 0.684 | 0.03 |
| Similarities | 0.697 | 0.668 | 0.029 |
| Digit Span | 0.692 | 0.681 | 0.011 |
| CVLT TL | 0.69 | 0.654 | 0.036 |
| Vocabulary | 0.653 | 0.633 | 0.02 |
| TMTB | 0.648 | 0.669 | -0.021 |
| BNT | 0.606 | 0.547 | 0.059 |
| Coding | 0.596 | 0.627 | -0.031 |
| Logical Memory 2 | 0.588 | 0.567 | 0.021 |
| Semantic fluency | 0.578 | 0.467 | 0.111 |
| Logical Memory 1 | 0.571 | 0.534 | 0.037 |
| TMTA | 0.554 | 0.572 | -0.018 |
| Pegboard R | 0.502 | 0.58 | -0.078 |
| Matrix Reasoning | 0.498 | 0.568 | -0.07 |
| Block Design | 0.493 | 0.541 | -0.048 |
| Pegboard L | 0.43 | 0.493 | -0.063 |
| ROCF Copy | 0.418 | 0.484 | -0.066 |
| WCST PR | 0.288 | 0.346 | -0.058 |
| WCST CC | 0.277 | 0.333 | -0.056 |

**Supplementary Table 6. Chi-square test comparison subtypes on cluster subclass for non-normative ICNs (Figure 5a)**

|  | ***n* Intact**  **(%)** | ***n* impaired**  **(%)** | ***n***  **HC**  **(%)** | **Impaired vs. Intact** | | | **Intact vs. HC** | | | **Impaired vs. HC** | | |
| --- | --- | --- | --- | --- | --- | --- | --- | --- | --- | --- | --- | --- |
|  |  |  |  | **Chi-square** | ***p*** | ***p* FDR** | **Chi-square** | ***p*** | ***p* FDR** | **Chi-square** | ***p*** | ***p* FDR** |
| Cluster 1 | 0.1905 | 0.3333 | 0.1087 | 2.0034 | 0.1569 | 0.6986 | 2.7405 | 0.0978 | 0.5486 | 8.4408 | 0.0037 | 0.0367 |
| Cluster 2 | 0.6667 | 0.5833 | 0.5543 | -0.5262 | 0.4682 | 0.7104 | 2.3522 | 0.1251 | 0.5486 | 0.1222 | 0.7266 | 0.9213 |
| Cluster 3 | 0.1270 | 0.0833 | 0.1087 | -0.3255 | 0.5683 | 0.7104 | 0.3256 | 0.5683 | 0.7103 | -0.0466 | 0.8291 | 0.9213 |
| Cluster 4 | 0.0794 | 0.1250 | 0.0543 | 0.4335 | 0.5103 | 0.7104 | 0.8805 | 0.3481 | 0.6735 | 2.2309 | 0.1353 | 0.3865 |
| Cluster 5 | 0.8095 | 0.9583 | 0.8370 | 3.0280 | 0.0818 | 0.6986 | -0.0693 | 0.7923 | 0.7923 | 2.6616 | 0.1028 | 0.3833 |
| Cluster 6 | 0.2222 | 0.3333 | 0.1522 | 1.1357 | 0.2866 | 0.6986 | 1.7021 | 0.1920 | 0.5486 | 4.7342 | 0.0296 | 0.1971 |
| Cluster 7 | 0.2381 | 0.3333 | 0.2717 | 0.8106 | 0.3680 | 0.6986 | -0.1030 | 0.7483 | 0.7923 | 0.5003 | 0.4793 | 0.8120 |
| Cluster 8 | 0.0476 | 0.0417 | 0.0326 | -0.0140 | 0.9057 | 0.9057 | 0.8023 | 0.3704 | 0.6735 | 0.3000 | 0.5839 | 0.8983 |
| Cluster 9 | 0.1429 | 0.2500 | 0.1957 | 1.3982 | 0.2370 | 0.6986 | -0.4708 | 0.4926 | 0.7103 | 0.5093 | 0.4754 | 0.8120 |
| Cluster 10 | 0.7302 | 0.7917 | 0.6957 | 0.3480 | 0.5552 | 0.7104 | 0.3689 | 0.5436 | 0.7103 | 1.0495 | 0.3056 | 0.7640 |
| Cluster 11 | 0.0952 | 0.1250 | 0.1196 | 0.1660 | 0.6837 | 0.7197 | -0.0732 | 0.7868 | 0.7923 | 0.0508 | 0.8216 | 0.9213 |
| Cluster 12 | 0.2698 | 0.4167 | 0.2500 | 1.7505 | 0.1858 | 0.6986 | 0.1873 | 0.6652 | 0.7826 | 3.0033 | 0.0831 | 0.3833 |
| Cluster 13 | 0.6825 | 0.5833 | 0.5543 | -0.7571 | 0.3842 | 0.6986 | 3.0130 | 0.0826 | 0.5486 | 0.1222 | 0.7266 | 0.9213 |
| Cluster 14 | 0.1270 | 0.2917 | 0.0761 | 3.3033 | 0.0691 | 0.6986 | 1.7363 | 0.1876 | 0.5486 | 9.8089 | 0.0017 | 0.0347 |
| Cluster 15 | 0.3175 | 0.3750 | 0.4130 | 0.2589 | 0.6109 | 0.7187 | -1.1542 | 0.2827 | 0.6292 | -0.0587 | 0.8085 | 0.9213 |
| Cluster 16 | 0.4444 | 0.3750 | 0.3696 | -0.3429 | 0.5582 | 0.7104 | 1.1520 | 0.2831 | 0.6292 | 0.0219 | 0.8823 | 0.9234 |
| Cluster 17 | 0.5079 | 0.4583 | 0.3913 | -0.1711 | 0.6792 | 0.7197 | 2.4770 | 0.1155 | 0.5486 | 0.4827 | 0.4872 | 0.8120 |
| Cluster 18 | 0.2381 | 0.3750 | 0.3043 | 1.6306 | 0.2016 | 0.6986 | -0.5806 | 0.4461 | 0.7103 | 0.5910 | 0.4420 | 0.8120 |
| Cluster 19 | 0.3175 | 0.2083 | 0.2283 | -1.0107 | 0.3147 | 0.6986 | 1.9557 | 0.1620 | 0.5486 | -0.0092 | 0.9234 | 0.9234 |
| Cluster 20 | 0.2222 | 0.3333 | 0.1957 | 1.1357 | 0.2866 | 0.6986 | 0.3276 | 0.5671 | 0.7103 | 2.4844 | 0.1150 | 0.3833 |

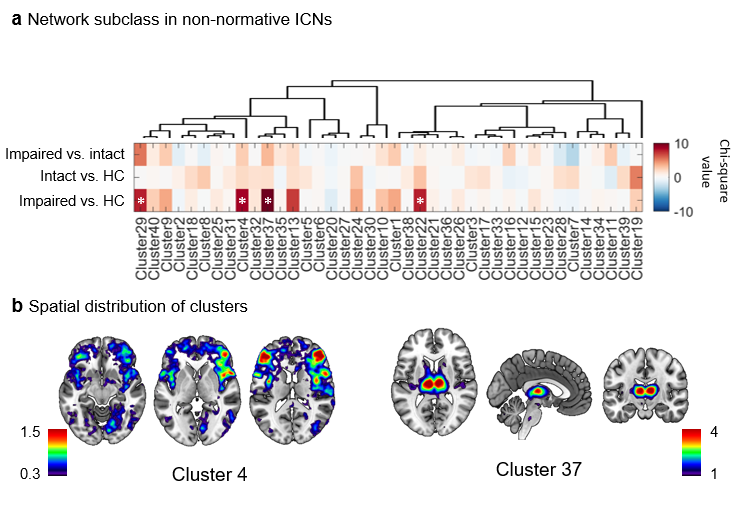

**Supplementary Figure 8. Forty network subclasses in non-normative ICNs.** To show the stability of the analysis in terms of the number of cluster subclasses, we repeated the main results in Figure 5 using 40 subclasses. **(a)** Agglomerative hierarchical clustering partitioned non-normative ICNs into 40 subclasses. Pseudo-colored plots showing the chi-square value of differences in the frequency of occurrence of all subclasses across our participant groups. Chi-square tests showed cognitively impaired patients owned more ICNs in the Cluster 4 and Cluster 37 subclasses. The number of ICNs in Cluster 29 and 22 was too small (less than 10) to be considered statistically, scientifically reliable. **(b)** By mapping the spatial distribution of these two subclasses, we found that Cluster 4 is predominantly a language-related network, while Cluster 37 is a thalamus-related network.

### References

Abraham, A., Pedregosa, F., Eickenberg, M., Gervais, P., Mueller, A., Kossaifi, J., Gramfort, A., Thirion, B., & Varoquaux, G. (2014). Machine learning for neuroimaging with scikit-learn. *Front Neuroinform*, *8*, 14. <https://doi.org/10.3389/fninf.2014.00014>

Achard, S., & Bullmore, E. (2007). Efficiency and cost of economical brain functional networks. *PLoS Comput Biol*, *3*(2), e17. <https://doi.org/10.1371/journal.pcbi.0030017>

Barcelo, F., Sanz, M., Molina, V., & Rubia, F. J. (1997). The Wisconsin Card Sorting Test and the assessment of frontal function: a validation study with event-related potentials. *Neuropsychologia*, *35*(4), 399-408. <https://doi.org/10.1016/s0028-3932(96)00096-6>

Benton, A., Hamsher, K. d., & Sivan, A. (1994). Manual for the multilingual aphasia examination. *Iowa City, IA: AJA Associates*.

Bowie, C. R., & Harvey, P. D. (2006). Administration and interpretation of the Trail Making Test. *Nat Protoc*, *1*(5), 2277-2281. <https://doi.org/10.1038/nprot.2006.390>

Ciric, R., Rosen, A. F. G., Erus, G., Cieslak, M., Adebimpe, A., Cook, P. A., Bassett, D. S., Davatzikos, C., Wolf, D. H., & Satterthwaite, T. D. (2018). Mitigating head motion artifact in functional connectivity MRI. *Nat Protoc*, *13*(12), 2801-2826. <https://doi.org/10.1038/s41596-018-0065-y>

Cox, R. W., & Hyde, J. S. (1997). Software tools for analysis and visualization of fMRI data. *NMR Biomed*, *10*(4-5), 171-178. <https://doi.org/10.1002/(sici)1099-1492(199706/08)10:4/5><171::aid-nbm453>3.0.co;2-l

Dale, A. M., Fischl, B., & Sereno, M. I. (1999). Cortical surface-based analysis. I. Segmentation and surface reconstruction. *Neuroimage*, *9*(2), 179-194. <https://doi.org/10.1006/nimg.1998.0395>

Delis, D. C. (2000). *California Verbal Learning Test, Second Edition: CvLT-II ; Adult Version ; Manual*. Pearson. <https://books.google.com/books?id=OfFlmwEACAAJ>

Esteban, O., Markiewicz, C. J., Blair, R. W., Moodie, C. A., Isik, A. I., Erramuzpe, A., Kent, J. D., Goncalves, M., DuPre, E., Snyder, M., Oya, H., Ghosh, S. S., Wright, J., Durnez, J., Poldrack, R. A., & Gorgolewski, K. J. (2019). fMRIPrep: a robust preprocessing pipeline for functional MRI. *Nat Methods*, *16*(1), 111-116. <https://doi.org/10.1038/s41592-018-0235-4>

Franz Petermann, -. (2008). *WAIS-IV : Wechsler Adult Intelligence Scale - Fourth Edition : Deutschsprachige Adaptation nach David Wechsler*. Pearson Assessment and Information. <https://books.google.com/books?id=s4j_mgEACAAJ>

Gladsjo, J. A., Schuman, C. C., Evans, J. D., Peavy, G. M., Miller, S. W., & Heaton, R. K. (1999). Norms for letter and category fluency: demographic corrections for age, education, and ethnicity. *Assessment*, *6*(2), 147-178. <https://doi.org/10.1177/107319119900600204>

Goodglass, H., Kaplan, E., & Weintraub, S. (2001). *BDAE: The Boston Diagnostic Aphasia Examination*. Lippincott Williams & Wilkins Philadelphia, PA.

Gorgolewski, K., Burns, C. D., Madison, C., Clark, D., Halchenko, Y. O., Waskom, M. L., & Ghosh, S. S. (2011). Nipype: a flexible, lightweight and extensible neuroimaging data processing framework in python. *Front Neuroinform*, *5*, 13. <https://doi.org/10.3389/fninf.2011.00013>

Jenkinson, M., Bannister, P., Brady, M., & Smith, S. (2002). Improved optimization for the robust and accurate linear registration and motion correction of brain images. *Neuroimage*, *17*(2), 825-841. <https://doi.org/10.1016/s1053-8119(02)91132-8>

Kim, J., Criaud, M., Cho, S. S., Diez-Cirarda, M., Mihaescu, A., Coakeley, S., Ghadery, C., Valli, M., Jacobs, M. F., Houle, S., & Strafella, A. P. (2017). Abnormal intrinsic brain functional network dynamics in Parkinson's disease. *Brain*, *140*(11), 2955-2967. <https://doi.org/10.1093/brain/awx233>

Latora, V., & Marchiori, M. (2001). Efficient behavior of small-world networks. *Phys Rev Lett*, *87*(19), 198701. <https://doi.org/10.1103/PhysRevLett.87.198701>

Power, J. D., Mitra, A., Laumann, T. O., Snyder, A. Z., Schlaggar, B. L., & Petersen, S. E. (2014). Methods to detect, characterize, and remove motion artifact in resting state fMRI. *Neuroimage*, *84*, 320-341. <https://doi.org/10.1016/j.neuroimage.2013.08.048>

Pruim, R. H. R., Mennes, M., van Rooij, D., Llera, A., Buitelaar, J. K., & Beckmann, C. F. (2015). ICA-AROMA: A robust ICA-based strategy for removing motion artifacts from fMRI data. *Neuroimage*, *112*, 267-277. <https://doi.org/10.1016/j.neuroimage.2015.02.064>

Rubinov, M., & Sporns, O. (2010). Complex network measures of brain connectivity: uses and interpretations. *Neuroimage*, *52*(3), 1059-1069. <https://doi.org/10.1016/j.neuroimage.2009.10.003>

Satterthwaite, T. D., Elliott, M. A., Gerraty, R. T., Ruparel, K., Loughead, J., Calkins, M. E., Eickhoff, S. B., Hakonarson, H., Gur, R. C., Gur, R. E., & Wolf, D. H. (2013a). An improved framework for confound regression and filtering for control of motion artifact in the preprocessing of resting-state functional connectivity data. *Neuroimage*, *64*, 240-256. <https://doi.org/10.1016/j.neuroimage.2012.08.052>

Satterthwaite, T. D., Elliott, M. A., Gerraty, R. T., Ruparel, K., Loughead, J., Calkins, M. E., Eickhoff, S. B., Hakonarson, H., Gur, R. C., Gur, R. E., & Wolf, D. H. (2013b). An improved framework for confound regression and filtering for control of motion artifact in the preprocessing of resting-state functional connectivity data [Article]. *Neuroimage*, *64*, 240-256. <https://doi.org/10.1016/j.neuroimage.2012.08.052>

Shin, M. S., Park, S. Y., Park, S. R., Seol, S. H., & Kwon, J. S. (2006). Clinical and empirical applications of the Rey-Osterrieth Complex Figure Test. *Nat Protoc*, *1*(2), 892-899. <https://doi.org/10.1038/nprot.2006.115>

Skogan, A. H., Oerbeck, B., Christiansen, C., Lande, H. L., & Egeland, J. (2018). Updated developmental norms for fine motor functions as measured by finger tapping speed and the Grooved Pegboard Test. *Dev Neuropsychol*, *43*(7), 551-565. <https://doi.org/10.1080/87565641.2018.1495724>

Smith, S. M., Fox, P. T., Miller, K. L., Glahn, D. C., Fox, P. M., Mackay, C. E., Filippini, N., Watkins, K. E., Toro, R., Laird, A. R., & Beckmann, C. F. (2009). Correspondence of the brain's functional architecture during activation and rest. *Proc Natl Acad Sci U S A*, *106*(31), 13040-13045. <https://doi.org/10.1073/pnas.0905267106>

Treiber, J. M., White, N. S., Steed, T. C., Bartsch, H., Holland, D., Farid, N., McDonald, C. R., Carter, B. S., Dale, A. M., & Chen, C. C. (2016). Characterization and Correction of Geometric Distortions in 814 Diffusion Weighted Images. *PLoS One*, *11*(3), e0152472. <https://doi.org/10.1371/journal.pone.0152472>

Tustison, N. J., Avants, B. B., Cook, P. A., Zheng, Y., Egan, A., Yushkevich, P. A., & Gee, J. C. (2010). N4ITK: improved N3 bias correction. *IEEE Trans Med Imaging*, *29*(6), 1310-1320. <https://doi.org/10.1109/TMI.2010.2046908>

Wang, S., Peterson, D. J., Gatenby, J. C., Li, W., Grabowski, T. J., & Madhyastha, T. M. (2017). Evaluation of Field Map and Nonlinear Registration Methods for Correction of Susceptibility Artifacts in Diffusion MRI. *Front Neuroinform*, *11*, 17. <https://doi.org/10.3389/fninf.2017.00017>

WECHSLER, D. (2022). *WMS-IV: WECHSLER MEMORY SCALE*. PEARSON. <https://books.google.com/books?id=XhyBzwEACAAJ>

Yeo, B. T., Krienen, F. M., Sepulcre, J., Sabuncu, M. R., Lashkari, D., Hollinshead, M., Roffman, J. L., Smoller, J. W., Zollei, L., Polimeni, J. R., Fischl, B., Liu, H., & Buckner, R. L. (2011). The organization of the human cerebral cortex estimated by intrinsic functional connectivity. *J Neurophysiol*, *106*(3), 1125-1165. <https://doi.org/10.1152/jn.00338.2011>

Zhang, Y., Brady, M., & Smith, S. (2001). Segmentation of brain MR images through a hidden Markov random field model and the expectation-maximization algorithm. *IEEE Trans Med Imaging*, *20*(1), 45-57. <https://doi.org/10.1109/42.906424>
